# Single-nucleus Multiomic Analyses Identifies Gene Regulatory Dynamics of Phenotypic Modulation in Human Aneurysmal Aortic Root

**DOI:** 10.1101/2024.02.27.582442

**Authors:** Xuanyu Liu, Qingyi Zeng, Hang Yang, Wenke Li, Qianlong Chen, Kunlun Yin, Zihang Pan, Kai Wang, Mingyao Luo, Chang Shu, Zhou Zhou

## Abstract

Aortic root aneurysm is a potentially life-threatening condition that may lead to aortic rupture and is often associated with genetic syndromes, such as Marfan syndrome (MFS). Although studies with MFS animal models have provided valuable insights into the pathogenesis of aortic root aneurysms, our understanding of the transcriptomic and epigenomic landscape in human aortic root tissue remains incomplete. This knowledge gap has impeded the development of effective targeted therapies. Here, this study performs the first integrative analysis of single-nucleus multiomic (gene expression and chromatin accessibility) and spatial transcriptomic sequencing data of human aortic root tissue under healthy and MFS conditions. Cell-type-specific transcriptomic and cis-regulatory profiles in the human aortic root are identified. Regulatory and spatial dynamics during phenotypic modulation of vascular smooth muscle cells (VSMCs), the cardinal cell type, are delineated. Moreover, candidate key regulators driving the phenotypic modulation of VSMC, such as *FOXN3*, *TEAD1*, *BACH2*, and *BACH1*, are identified. *In vitro* experiments demonstrate that FOXN3 functions as a novel key regulator for maintaining the contractile phenotype of human aortic VSMCs through targeting ACTA2. These findings provide novel insights into the regulatory and spatial dynamics during phenotypic modulation in the aneurysmal aortic root of humans.

## 1. Introduction

Aortic root aneurysm begins as an asymptomatic dilatation of the aortic root, enlarges progressively over time, and may ultimately lead to aortic dissection or rupture^[1]^. Many cases of aortic root aneurysms are associated with genetic syndromes such as Marfan syndrome (MFS)^[2]^. MFS is a highly penetrant connective tissue disorder (caused by mutations in *FBN1* encoding fibrillin-1) with pleiotropic manifestations in the ocular, skeletal, and cardiovascular systems^[1]^. The cardinal and potentially life-threatening manifestation of MFS is aortic root aneurysm^[3]^. Medications (e.g., β-adrenergic receptor blockers) to slow aortic growth or prophylactic aortic surgery to prevent dissections can improve the lifespan of MFS patients^[4]^. However, effective targeted medical therapies for aortic root aneurysms are still limited, partially due to the incomplete understanding of the regulatory mechanisms underlying the pathophysiological changes. While extensive research has focused on the roles of the TGFβ and angiotensin II signaling pathways in the pathogenesis of aortic root aneurysms^[5]^, the gene regulatory programs driving the pathological changes in the aneurysmal aortic root of human MFS patients remain elusive.

To date, mechanistic insights into the pathogenesis of aortic root aneurysms have stemmed mostly from animal models, e.g., *Fbn1*^C1039G/+^ mice. Nevertheless, MFS mouse models may not accurately mimic the pathophysiological changes that occur in humans. For example, the MFS mouse models develop aortic aneurysms, but the aneurysms rarely progress to dissection or rupture^[5]^. These discrepancies partially explain why therapeutic efficacy in mice does not accurately predict clinical trial success in patients, and underscore the importance of delineating the pathophysiological changes in tissues derived from human patients.

The pathogenesis of aortic root aneurysms involves multiple cell types that undergo complex phenotypic modulation. Vascular smooth muscle cells (VSMCs) are the cardinal cell type of the aortic wall with remarkable plasticity. Phenotypic modulation of VSMCs from a differentiated contractile phenotype toward a dedifferentiated synthetic phenotype with increased proliferation and migration underlies the pathogenesis of many vascular diseases such as aortic aneurysms and atherosclerosis^[6]^. In MFS mice, the transcriptomic dynamics during the phenotypic modulation of aortic root/ascending aneurysm tissue have been delineated using single-cell RNA sequencing^[7]^. Cell-type-specific transcriptomic changes have been uncovered in human ascending aortic aneurysms with a single-cell/nucleus RNA sequencing approach^[8,9]^. However, we still lack comprehensive knowledge concerning the alterations in the aneurysmal aortic root tissues of human MFS at the single-cell/nucleus level. In addition, the phenotypic heterogeneity of VSMCs is expected to be associated with spatial locations within the aortic wall. An integrative analysis of single-cell/nucleus and spatial transcriptomic datasets would unravel both transcriptomic and spatial dynamics of phenotype modulation in the aneurysmal aortic root.

Gene expression is governed by cis-regulatory elements (CREs) in a spatiotemporal and cell-type-specific manner, including enhancers, promoters, and insulators^[10]^. A detailed atlas of cell-type-specific accessible CREs in tissues under healthy and diseased conditions is fundamental for delineating regulatory mechanisms underlying pathogenesis. In addition, single-cell epigenomic data facilitate the interpretation of the growing number of aortic disease-associated genetic loci identified by genome-wide association studies, since the majority of the risk variants reside in noncoding regions^[11]^. Cell-type-specific candidate CREs (cCREs) have been uncovered in multiple human tissue types with the single-cell assay for transposase-accessible chromatin using sequencing (scATAC-seq), such as the human coronary artery^[12]^ and myocardium^[13]^. However, such an atlas is still lacking for the human aortic root. Recent technical advances have enabled joint profiling of gene expression and chromatin accessibility from the same nucleus (represented by 10X Genomics Chromium Single Cell Multiome ATAC + Gene Expression), which should greatly increase the power for delineating gene regulatory mechanisms.

In this study, through an integrative analysis of single-nucleus multiomic (gene expression and chromatin accessibility) and spatial transcriptomic sequencing data of human aortic root tissues under healthy and MFS conditions, we built the first atlas of gene expression and chromatin accessibility in the human aortic root at single-nucleus and spatial resolution. The unique dataset provided novel insights into the regulatory and spatial dynamics during the phenotypic modulation of VSMCs in the aneurysmal aortic root. FOXN3 was identified as a novel key regulator for maintaining the contractile phenotype of human aortic VSMCs, which may serve as a potential therapeutic target.

## 2. Results

### 2.1. Single-nucleus multiomic and spatial transcriptomic sequencing of aortic root tissues from MFS patients and healthy controls

Aneurysmal aortic root tissues from MFS patients (n = 6) were collected during surgery. As controls, normal aortic root tissues from heart transplantation recipients (n = 6) were also collected. The control group (CTRL) was ethnicity-(Chinese) and age-matched (MFS: 35.8 ± 6.1 years old; CTRL: 41.8 ±6.1 years old, *P* = 0.25, Wilcoxon rank sum test, two-tailed) with the MFS group. Detailed demographic and clinical information of the enrolled subjects is outlined in Table S1, Supporting Information. The staining of aortic root tissue sections from individuals with Marfan syndrome (MFS) using hematoxylin and eosin (H&E) and Elastic van Gieson (EVG) revealed several histopathological hallmarks of aortic aneurysms. Specifically, the samples exhibited medial layer degeneration, which was characterized by the fragmentation or loss of elastic fibers, the accumulation of mucoid extracellular matrix, and the loss of smooth muscle nuclei (Figure S1, Supporting Information). Samples from four subjects of each group were individually subjected to single-nucleus multiomic sequencing using the protocol of 10X Genomics Chromium Single Cell Multiome ATAC + Gene Expression, which enables joint profiling of gene expression and chromatin accessibility from the same nucleus (Figure 1A). After stringent quality control (Figures S2 and S3; Table S2, Supporting Information), a total of 36,316 nuclei with 25,887 expressed genes and 137,311 accessible cCREs were obtained. The identified cCREs covered 2.8% of the human genome, 75.1% of which overlapped previously documented cCREs in ENCODE (926,535 cCREs in various human tissues and cells). Only 13.8% of the identified cCREs were annotated within promoter regions (Figure S4, Supporting Information). To further infer the regulatory roles of the cCREs, 12.3% of them were linked to genes by computing the correlation between their accessibility and expression of the nearby genes (Table S3, Supporting Information). In addition, to study gene expression in a spatial context, spatial transcriptomic assays (10X Visium Spatial Gene Expression) were applied to aortic root tissue sections from two subjects of each group. In total, 1,651 to 3,036 spots were detected to be over tissue (Table S2, Supporting Information). A web-based interactive interface (http://multiomeMFS.fwgenetics.org/) was established for all the datasets.

**Figure 1.**
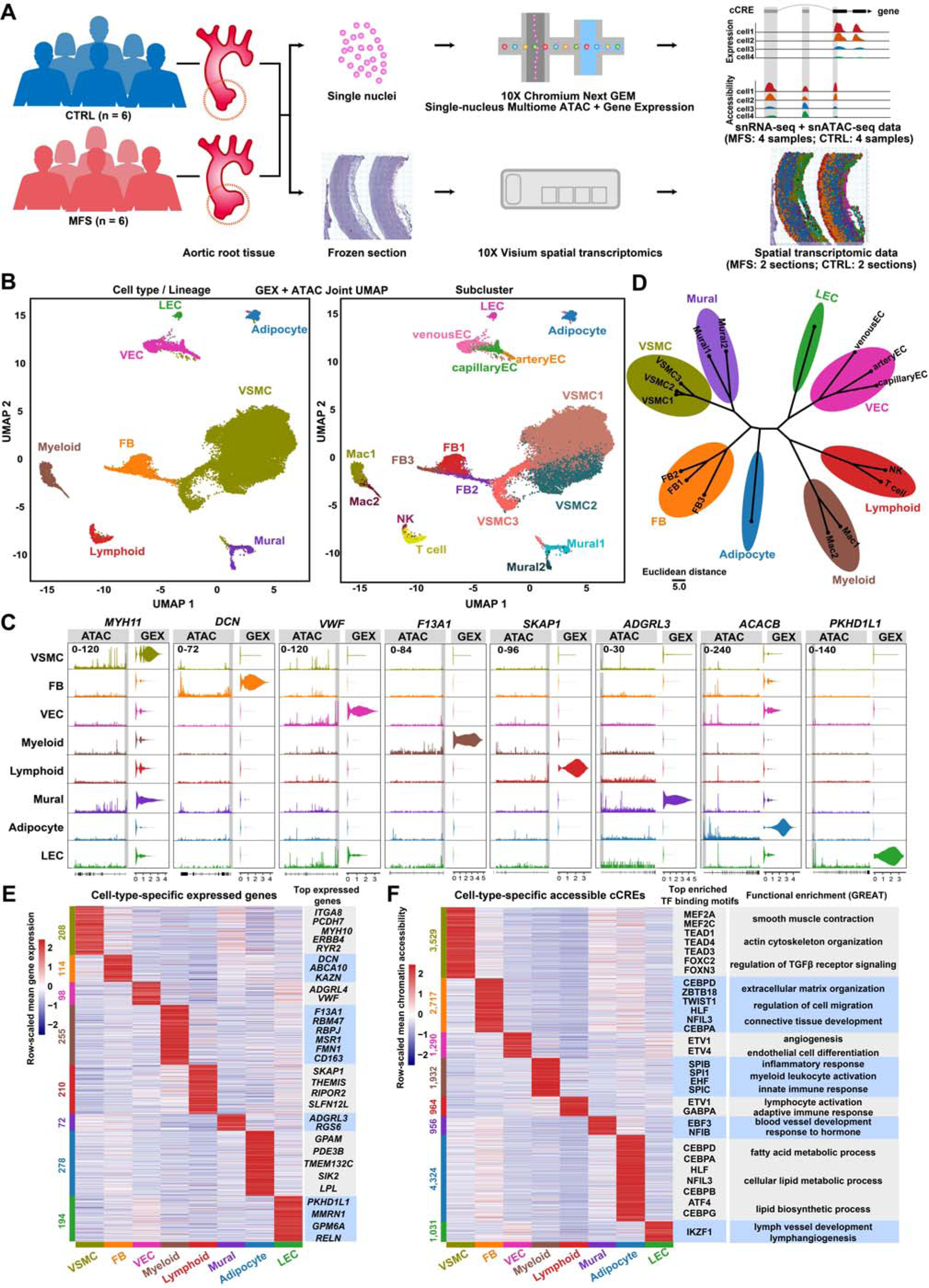
Single-nucleus multiomic analysis reveals cell-type-specific expressed genes and accessible cCREs in aortic root tissues from MFS patients and healthy controls. A) Schematic representation of the procedure for generating the sequencing data. Aneurysmal aortic root tissues from MFS patients (n = 6) and normal aortic root tissues from heart transplantation recipients (n = 6) were collected. Samples from four subjects (two males and two females) of each group were individually subjected to single-nucleus multiomic sequencing. Tissue sections from two subjects (one male and one female) of each group were subjected to spatial transcriptomic assays. MFS: Marfan syndrome; CTRL: control. B) Joint UMAP visualization of cell types and subclusters that represents the measurements of both gene expression and chromatin accessibility modalities. GEX: gene expression; ATAC: assay for transposase-accessible chromatin. C) Gene expression and chromatin accessibility profiles for marker genes of each cell type/lineage. The promoter region is highlighted in gray. D) Hierarchical clustering of all the subclusters. The top 30 PCA components of the gene expression data and the 2-30 integrated LSI components of the chromatin accessibility data were considered. E) Heatmap showing the expression of cell-type-specific expressed genes. The significance threshold for gene expression was set to a log2(fold change) value > 0.5 and a *p*-value adjusted for multiple testing < 0.05 (likelihood-ratio test). F) Heatmap showing the accessibility of cell-type-specific assessable cCREs. The significance threshold for differential accessibility was set to a log2(fold change) value > 0.25 and a *p*-value adjusted for multiple testing < 0.01 (logistic regression test). Top TF binding motifs and representative Gene Ontology terms (inferred by the tool GREAT) enriched for each cell type are shown. The significance threshold was set to a Bonferroni-corrected *p*-value < 0.05 (hypergeometric test). The cell-type-specific expressed genes or assessable cCREs were detected using the function “FindAllMarkers” of the Seurat package. FB: fibroblast; LEC: lymphatic endothelial cell; VEC: vascular endothelial cell; VSMC: vascular smooth muscle cell.

### 2.2. Cell-type-specific transcriptomic and cis-regulatory profiles in human aortic root tissues under healthy and MFS conditions

Unsupervised clustering of all the nuclei enabled the identification of canonical cell types in the aorta encompassing VSMCs, fibroblasts, mural cells, vascular endothelial cells, lymphatic endothelial cells, adipocytes, myeloid cells, and lymphoid cells (Figure 1B), which was supported by the expression and promoter accessibility of established markers (Figure 1C). Notably, VSMCs accounted for 76.9% of all the nuclei, which is consistent with the expected cellular composition of the human aorta^[9,14]^. Subclustering of the major cell types further revealed within-lineage subclusters, e.g., three subclusters for the VSMC lineage (Figure 1B). These subclusters/cell types could be distinguished in the UMAP embeddings based on joint datasets (gene expression + chromatin accessibility) or a separate dataset (Figure S5, Supporting Information). Furthermore, the result of hierarchical clustering supported the expected degree of similarity among clusters, reflecting the robustness of the clustering (Figure 1D). Leveraging the single-nucleus multiomic data, we identified both cell-type-specific gene expression (Figure 1E; Table S4, Supporting Information) and chromatin accessibility (Figure 1F; Table S5, Supporting Information) in the human aortic root. Then, transcription factor (TF) binding motifs enriched in the cCREs specifically accessible in each lineage were identified (Figure 1F and Table S6, Supporting Information). For example, VSMC-specific assessable cCREs were enriched for TF motifs such as MEF2 (Myocyte Enhancer Factor 2), TEAD (transcriptionally enhanced associate domain), and forkhead box TF families. Functional enrichment of the cell-type-specific accessible cCREs was inferred by using the Genomic Regions Enrichment of Annotations Tool (GREAT)^[15]^. For example, VSMC-specific accessible cCREs were enriched for VSMC-related terms, such as smooth muscle contraction and regulation of TGFβ receptor signaling (Figure 1F). In addition, subcluster-specific gene expression and chromatin accessibility were also identified (Figure S6, Supporting Information). Notably, VSMC1-specific assessable cCREs were enriched for the motif of SRF (serum response factor), a TF regulator with a central role in regulating the expression of smooth muscle-specific contractile genes^[16]^.

### 2.3. Cellular compositional alterations and cell-type-resolved regulatory changes in MFS compared with the controls

The two groups showed a high degree of overall similarity in terms of cell-type composition, as shown in Figure 2A (also see Figures S7 and S8, Supporting Information). This finding is consistent with a previous single-nucleus study on ascending aortic aneurysums^[9]^. Nevertheless, VSMC2, a VSMC subcluster, exhibited significant expansion in MFS samples compared with the controls (Figure 2B; a Bayesian method implemented in scCODA), reflecting its association with MFS. The expression signature of VSMC2 (high levels of *COL8A1* and *SERPINE1*)^[8]^ suggested the identity of modulated VSMCs (Table S4, Supporting Information). In addition, our data revealed that a considerable proportion (20∼30%) of modulated VSMCs were present in both healthy and aneurysmal aortic root tissues of humans, unlike the observation in MFS mouse models (few modulated VSMCs in the healthy condition)^[7]^.

**Figure 2.**
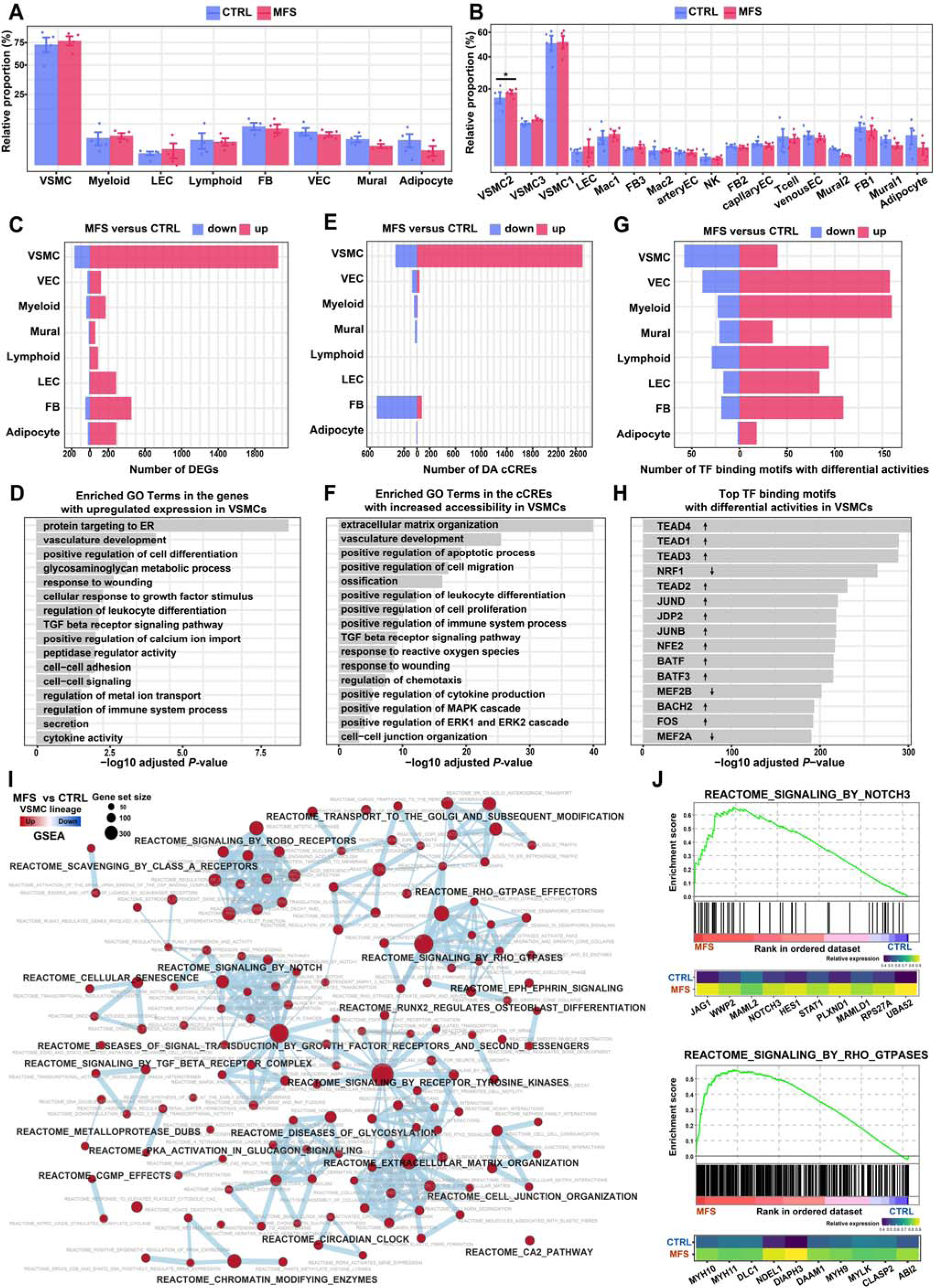
Cellular compositional alterations and cell-type-specific regulatory changes in MFS versus CTRL samples revealed by single-nucleus multiomic analysis. A) Relative proportion of each cell type in the aortic root tissues from the MFS and CTRL groups. The data on the y-axis were square-root transformed for better visualization. B) Relative proportion of each subcluster in the aortic root tissues from MFS or CTRL. In A and B, *: statistically significant change. Differential compositional testing was performed using a Bayesian approach implemented in scCODA. The data are presented as the mean ± SEM (n=4 for each group). C) Number of differentially expressed genes (DEGs) in each cell type between the MFS and CTRL groups. The significance threshold was set to an absolute log2 fold change >1 and a Bonferroni-adjusted *p*-value < 0.05. The statistical method implemented in DEsingle was used. D) Representative GO terms enriched in the upregulated genes in VSMCs from MFS versus CTRL. Bonferroni-corrected *p*-value < 0.05. The hypergeometric tests implemented in ClueGO were used. E) Number of differentially accessible (DA) cCREs in each cell type between the MFS and CTRL groups. The significance threshold was set to a Bonferroni-adjusted *p*-value < 0.05 and an absolute of log2 fold change > 0.1. The logistic regression test implemented in Seurat was used. F) Representative GO terms enriched in the cCREs with increased accessibility in VSMCs from MFS versus CTRL. Bonferroni-corrected *p*-value < 0.05. The hypergeometric test implemented in GREAT was used. G) Number of TF binding motifs with differential activities in each cell type between MFS and CTRL. Wilcoxon rank-sum test, two-tailed, FDR < 0.05. H) Top TF binding motifs with differential activities in the VSMCs from MFS patients versus the CTRL group. Up arrow: increased activity. Down arrow: decreased activity. I) Network view of the dysregulated REACTOME pathways in VSMCs from MFS patients versus the CTRL group inferred by gene set enrichment analysis (GSEA). An FDR < 0.05 was considered to indicate statistical significance. J) Enrichment plots (upper panel) for representative signaling pathways upregulated in the VSMCs of MFS and heatmaps showing the average expression of leading-edge genes in each condition (lower panel). The vertical lines in the enrichment plot show where the members of the gene set appear in the ranked list of genes. Leading-edge genes: the subset of genes in the gene set that contribute most to the enrichment. FB: fibroblast; LEC: lymphatic endothelial cell; NES: normalized enrichment score. VEC: vascular endothelial cell; VSMC: vascular smooth muscle cell.

Despite the absence of a large discrepancy in cellular composition between the groups, differentially expressed genes (DEGs; Figure 2C,D; Table S7, Supporting Information) and differentially accessible (DA) cCREs (Figure 2E,F; Table S8, Supporting Information) were detected between conditions, especially in VSMCs. The DEGs upregulated in VSMCs under the MFS condition were enriched for Gene Ontology (GO) terms related to VSMC dedifferentiation or phenotypic switching such as vasculature development, positive regulation of cell differentiation, extracellular matrix remodeling, and TGF beta receptor signaling pathway (Figure 2D). The downregulated genes in VSMCs under the MFS condition were enriched for pathways related to adrenergic receptor activity and smooth muscle contraction regulation (Figure S9a, Supporting Information). The functional enrichment of DA cCREs with altered accessibility in VSMCs under the condition of MFS was generally consistent with that of DEGs (Figure 2F; Figure S9b, Supporting Information). We also detected TF binding motifs with significantly altered activity (Figure 2G,H; Table S9, Supporting Information). Notably, the activity of TEAD binding motifs (TEAD1, TEAD2, TEAD3, and TEAD4) was significantly increased in MFS, consistent with the knowledge that TEAD TF family members function as regulators of VSMC phenotypic modulation^[16]^. To systematically detect dysregulated pathways in which the expression of genes changed in a small but concordant way, we performed gene set enrichment analysis (GSEA) and found upregulated signaling pathways in VSMCs under MFS conditions (Figure 2I; Table S10, Supporting Information), such as NOTCH3, Rho GTPases, ROBO receptor, TGF beta receptor complex, and receptor tyrosine kinase signaling pathways (Figure 2I,J).

### 2.4. Phenotypic spectrum and regulatory dynamics during the phenotypic modulation of VSMCs in the human aortic root

The single-nucleus dataset allowed us to examine the phenotypic spectrum of the highly plastic VSMCs, which encompassed three subclusters (Figure 3A-C; Table S11, Supporting Information). VSMC1 expressed high levels of *RYR2,* which encodes an intracellular Ca^2+^ release channel associated with the contractility of VSMCs^[17]^, and canonical contractile markers such as *ACTA2* and *CNN1*. These findings suggest that VSMC1 represents a contractile VSMC phenotype. VSMC2 expressed high levels of the signature genes of modulated VSMCs such as *COL8A1* and *SERPINE1*^[8]^. Notably, VSMC3 also expressed the signature genes of modulated VSMCs (e.g., *COL8A1*) but harbored a different expression profile from that of VSMC2 (e.g., high expression levels of *LAMA2* and *CFH*). Reanalysis of a publicly available single-cell dataset of aortic root tissue from an MFS patient also supported that VSMC3 represents a special state of the phenotypic spectrum of VSMCs rather than fibroblasts (Figure S10, Supporting Information). Pseudotime ordering of the VSMCs revealed that VSMC1 and VSMC3 may represent the two extremes of the phenotypic spectrum (Figure 3B). Through single-molecule fluorescent in situ hybridization (smFISH), we confirmed the presence of these two subclusters in human aortic root tissue, where they exhibited discrepancy in spatial distribution: *RYR2* ^high^ VSMC1 cells were mainly located in the media close to the adventitia, while *CFH* ^high^ VSMC3 cells were preferentially located in the media close to the intima (Figure 3D). As expected, the relative proportion of unmodulated contractile VSMC1 cells was decreased in MFS versus CTRL conditions, whereas the proportion of modulated VSMC2 cells was increased (Figure 3E; a Bayesian method implemented in scCODA). Plots of nucleus density also reflected altered transcriptomic states of VSMCs in MFS (Figure S11, Supporting Information).

**Figure 3.**
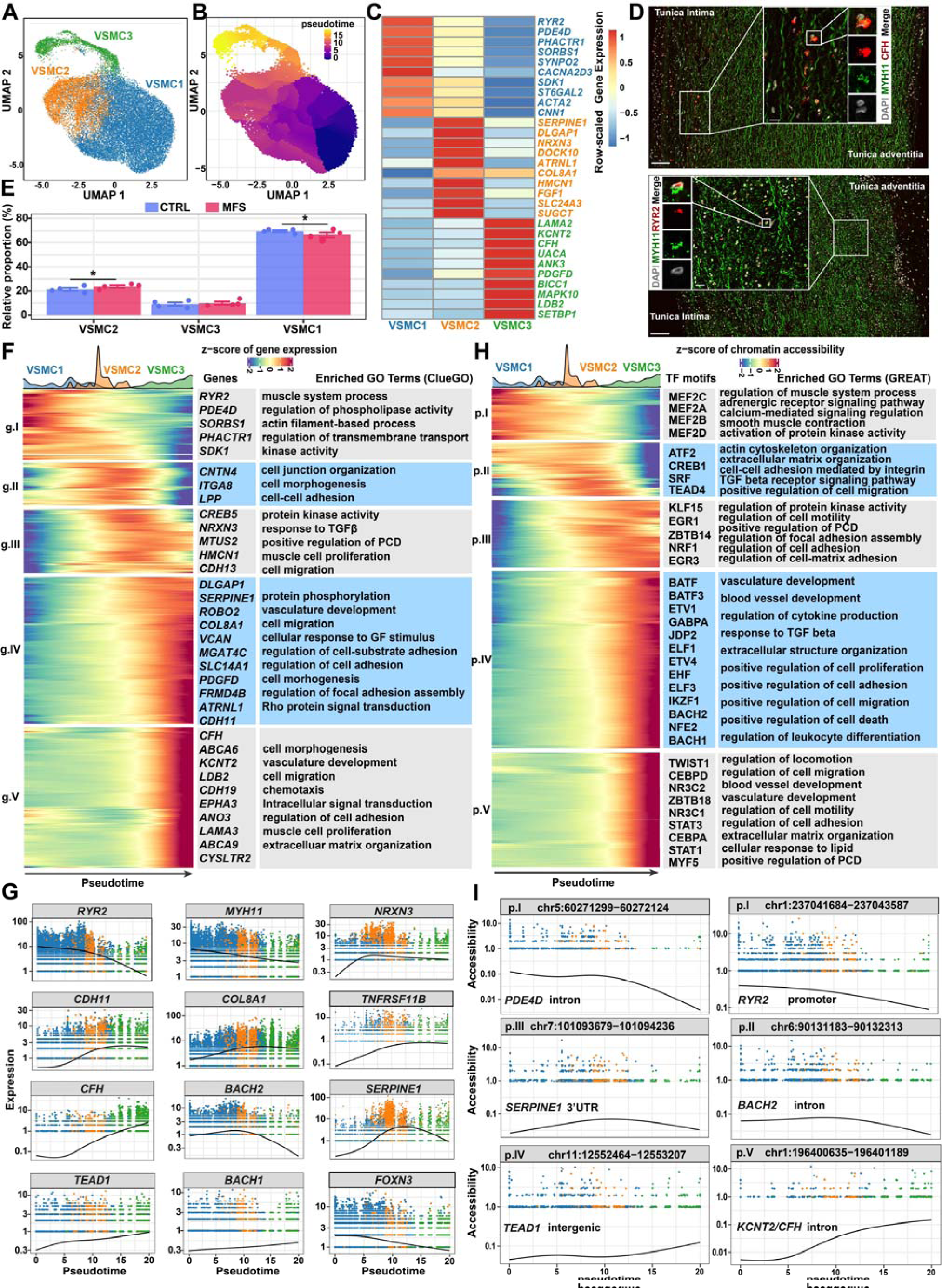
Phenotypic spectrum and regulatory dynamics during the phenotypic modulation of VSMCs in human aortic root tissue. A) UMAP plot showing the subclusters of VSMCs. B) UMAP plot showing the pseudotime inferred by Monocle3. C) Heatmap showing the expression of the top signature genes for each subcluster. D) smFISH confirmed the presence of *RYR2* ^high^ and *CFH* ^high^ VSMCs, the two extremes of the phenotypic spectrum of VSMCs. E) Relative proportion of each subcluster in VSMCs from each condition. The data are presented as the mean ± SEM (n=4 for each group). * statistically significant change. Differential compositional testing was conducted using a Bayesian approach implemented in scCODA. F) Heatmap showing the gene expression dynamics during the phenotypic modulation of VSMCs. Pseudotime ordering was performed using Monocle3. The significance threshold was set to a q-value < 0.05. Representative genes and enriched GO terms for each gene cluster are shown. G) Smoothed curves of representative genes whose expression changed as a function of pseudotime. H) Heatmap showing chromatin accessibility dynamics during the phenotypic modulation of VSMCs. The significance threshold was set to a q-value < 0.001. The top enriched TF motifs and GO terms for each peak cluster are shown. I) Smoothed curves of representative cCREs whose accessibility changed as a function of pseudotime. In F and G, the genes or cCREs that changed as a function of pseudotime were detected with graph-autocorrelation analysis by using the “graph_test” function in Monocle3.

Furthermore, by identifying genes or cCREs whose expression or accessibility changed as a function of pseudotime, the gene expression and chromatin accessibility dynamics during the phenotypic modulation of VSMCs were uncovered (Figure 3F,G; Table S12, Supporting Information). Functional enrichment analysis of these genes or cCREs suggested that the pseudotime ordering recapitulated the phenotypic modulation of VSMCs from a contractile phenotype toward a synthetic phenotype with increased proliferation and migration. The expression of markers for the contractile phenotype (e.g., *RYR2* and *MYH11*) gradually decreased over pseudotime, while markers for the modulated synthetic phenotype (e.g., *COL8A1*, *SERPINE1*, and *TNFRSF11B*) increased (Figure 3H). Notably, some TF genes, such as *FOXN3*, *BACH1*, *BACH2*, and *TEAD1*, were dynamically expressed over pseudotime, reflecting their potential role in phenotypic switching. Similarly, we also identified cCREs whose accessibility dynamically changed over pseudotime. For example, the accessibility of an intergenic cCRE (chr11:12552464-12553207) was identified to be positively correlated with the expression of *TEAD1*, suggesting a putative enhancer for this regulator (Figure 3I).

### 2.5. Candidate key regulators potentially driving the phenotypic modulation of VSMCs and pathogenesis of aortic root aneurysms

To identify potential key regulators driving the phenotypic modulation of VSMCs, we first performed single-cell weighted gene coexpression network analysis (scWGCNA). Nine gene coexpression modules in VSMCs were identified (Figure 4A; Table S13, Supporting Information). Two major modules whose expression was significantly changed in MFS were detected by comparing the module expression scores between conditions: module M7 had significantly lower expression scores in MFS versus CTRL conditions, while module M9 had significantly higher expression scores in MFS (Figure 4B; *p*-value < 0.05, Wilcoxon rank-sum test, two-tailed). The expression distribution in the UMAP projection showed that the VSMCs expressing module M7 generally overlapped with the contractile subcluster VSMC1 and that the VSMCs expressing module M9 generally overlapped with the modulated subclusters VSMC2 and VSMC3 (Figure 4C). In agreement with this, the top hub genes of M7 included markers for contractile VSMCs (e.g., *PDE4D* and *RYR2*), while the top hub genes of M9 included markers for modulated VSMCs (e.g., *CDH11*; Figure 4D). Notably, the core TFs of M7 and M9 included TFs that were dynamically expressed over pseudotime, such as *BACH2*, *FOXN3*, and *TEAD1* (Figure 4D). Overall, we identified gene coexpression modules associated with the contractile or modulated phenotypes of VSMCs.

**Figure 4.**
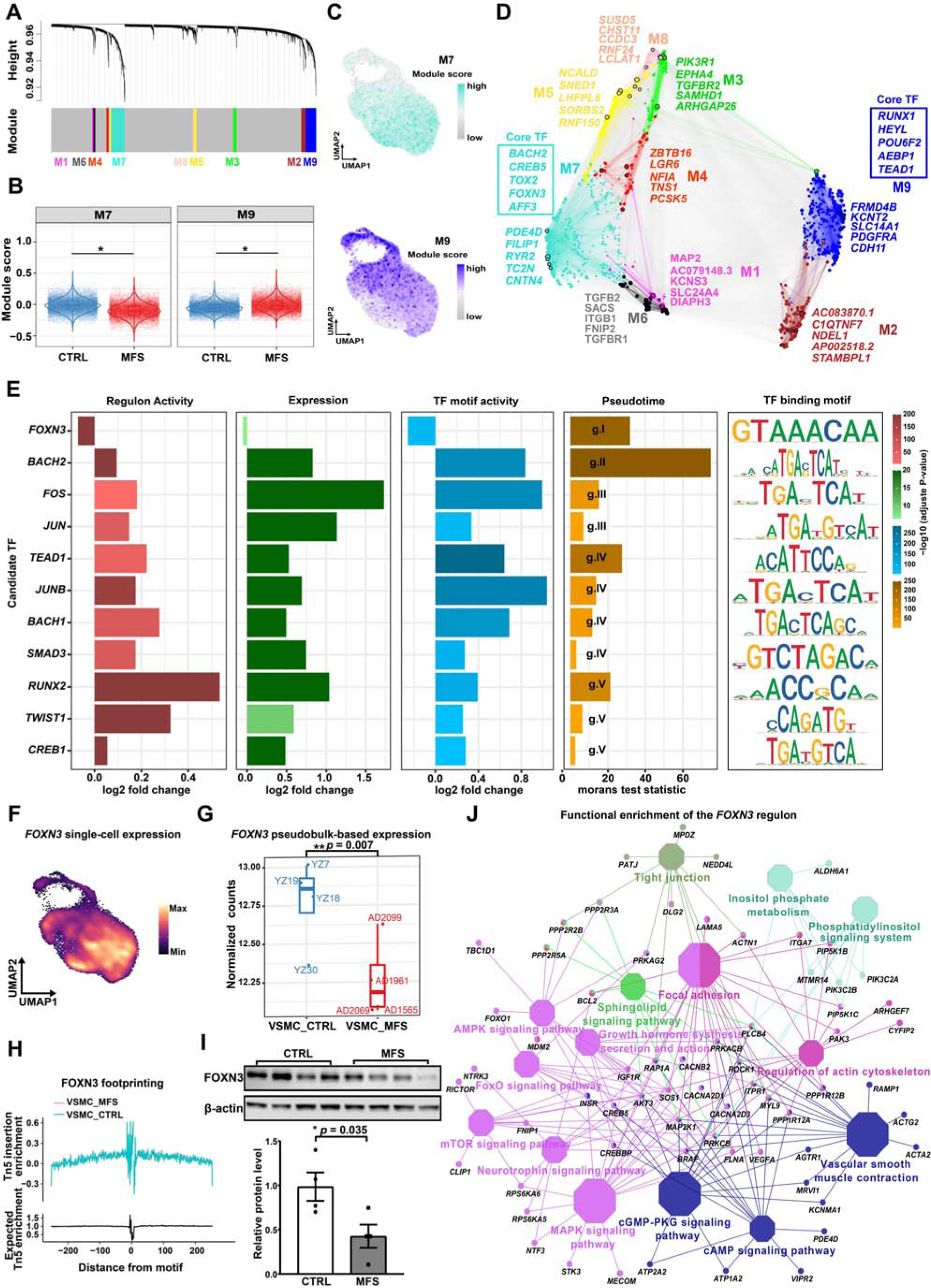
Candidate key regulators potentially driving the phenotypic modulation of VSMCs and the pathogenesis of aortic root aneurysms. A) Dendrogram showing the coexpression modules of the VSMCs identified by scWGCNA. B) VSMCs exhibited differential expression activities for the two largest modules, M7 and M9, in VSMCs of the MFS group versus the CTRL group. *: *p*-value < 0.05, Wilcoxon rank-sum test, two-tailed (CTRL: 14449 nuclei, MFS: 13497 nuclei). C) UMAP plots showing the expression distribution of modules M7 (upper panel) and M9 (lower panel) across all VSMCs. D) Gene regulatory network of VSMCs color-coded by co-expression modules. The top 5 hub genes of each module are shown. The core TFs of M7 and M9 inferred based on the centrality of the network are shown in boxes. E) Potentially key regulators involved in the phenotypic modulation of VSMCs supported by multiple layers of evidence including regulon activity, expression, TF motif activity, and pseudotime ordering. F) UMAP plot showing the distribution of single-nucleus expression of the TF *FOXN3*. The visualization was enhanced by using the R package Nebulosa to recover the signal from dropped-out features. G) Significantly downregulated pseudobulk expression of *FOXN3* in VSMCs from MFS patients versus CTRLs. **: *p*-value < 0.01, Wilcoxon rank-sum test, two-tailed (n = 4 for each group). H) TF footprinting differences of *FOXN3* between VSMCs from MFS patients and CTRLs. I) Western blot showing a significant decrease of the protein level of FOXN3 in the tunica media of the aortic root tissues from MFS patients compared to CTRLs. *: *p*-value < 0.05, Student’s t-test, two-tailed. The data are presented as the mean ± SEM (n = 4 for each group). J) Network plot showing the functional enrichment of the predicted *FOXN3* regulon (TF and its targets). The significance threshold was set to a Bonferroni-corrected *p*-value <0.05. The hypergeometric test implemented in ClueGO was used. Each octagon denotes an overrepresented REACTOME pathway. A larger size reflects a smaller adjusted *p*-value.

Next, we prioritized the potential key regulators driving the phenotypic modulation of VSMCs by integrating multiple pieces of evidence including significant differences in TF gene expression, regulon (a TF and its predicted targets) expression activity (Table S14, Supporting Information), and TF binding motif activity between conditions, as well as the pseudotime ordering results (Figure 4E). A total of 11 TFs were obtained, including TGFβ pathway-associated regulators (*SMAD3* and *TWIST1*), stress-responsive regulators (*FOS*, *JUN*, and *JUNB*), BACH family regulators (*BACH1* and *BACH2*), calcification (*RUNX2*), and others (*TEAD1*, *FOXN3,* and *CREB1*). TEAD1, a previously reported repressor of contractile gene expression in VSMCs^[16]^, was in the list of our candidate key regulators and supported by multiple pieces of evidence (Figure S12, Supporting Information). Notably, *FOXN3*, which was mainly expressed in contractile VSMCs (Figure 4F), was the only regulator whose expression and regulon activity were significantly decreased in MFS versus CTRL conditions; this finding was also supported by pseudobulk-based expression analysis (Figure 4G). Chromatin accessibility data showed stronger footprinting of *FOXN3* in VSMCs of the MFS group compared to those of the CTRL group. (Figure 4H). Western blot assays confirmed that the protein level of FOXN3 was significantly decreased in the media of the aortic root tissues from MFS patients (Figure 4I; *p*-valu*e* < 0.05). Functional enrichment of the FOXN3 regulon suggested that FOXN3 may target important pathways related to contractility, phenotypic modulation of VSMCs, and pathogenesis of aortic aneurysms, such as “vascular smooth muscle contraction”, “regulation of actin cytoskeleton”, and “focal adhesion” (Figure 4J). Together, our data uncovered candidate key regulators, for example, *FOXN3*, that potentially drive the phenotypic modulation of VSMCs and the pathogenesis of aortic root aneurysms.

### 2.6. Spatially resolved transcriptome showing the phenotypic spectrum of VSMCs across the tunica media of the human aortic root

Due to differences in factors such as mechanical loading and cellular microenvironment, VSMCs may exhibit phenotypic differences across the aortic wall. Spatial transcriptomic assays coupled with single-nucleus datasets offered us an opportunity to dissect the phenotypic spectrum of VSMCs across the tunica media. Through unsupervised clustering, nine spatial spot clusters were detected (Figure 5A,B). According to their expression signatures, the spot clusters were annotated as their major cell types (Figure 5C,D). Notably, the spot clusters annotated as VSMCs were located closely in the UMAP space and constituted a continuum of phenotypic changes from highly contractile VSMCs (SC4) to unmodulated contractile VSMCs (SC0) to modulated VSMCs (SC1). These spot clusters displayed a radial distribution across the media from the outer layer (the adventitial side) to the inner layer (the intimal side; Figure 5B). By using the label transfer workflow of Seurat, the single-nucleus data and spatial transcriptomic data were integrated, and the three VSMC subclusters were spatially confirmed in the tissue sections (Figure 5E). The contractile subcluster VSMC1 was located mainly in the outer layer of the tunica media, while the modulated subclusters VSMC2 and VSMC3 were located mainly in the inner layer. VSMC3 generally corresponded to spot cluster SC6. As expected, the relative proportion of spot cluster SC1 (modulated VSMCs) was increased in MFS samples compared with controls, while spot clusters SC0 and SC4 (unmodulated contractile VSMCs) were decreased (Figure 5F). The regulon activity of *FOXN3* was especially high in the outer layer of the tunica media whereas the regulon activity of *TEAD1* was especially high in the inner layer (Figure 5G), reflecting the association of these two regulators with different VSMC phenotypes. In analogy with pseudotime ordering, we performed pseudospace trajectory analysis of the VSMC spatial spots (SC4, SC0, and SC1; Figure 5H,I). This analysis allowed us to uncover the expression dynamics of VSMCs across the tunica media from the outer layer (unmodulated) to the inner layer (modulated; Figure 5J), which also represented the expression dynamics during the phenotypic modulation of VSMCs (Table S15, Supporting Information). The representative markers (e.g., *MYH11*) and candidate key regulators (e.g., *FOXN3*) were expressed dynamically along the pseudospace (Figure 5K), thus confirming the results of our single-nucleus data analyses. Overall, the spatially resolved transcriptome suggests that the modulated VSMCs are not only associated with the diseased states (i.e., increased relative proportion in MFS) but also associated with the spatial location (i.e., the inner layer of the tunica media).

**Figure 5.**
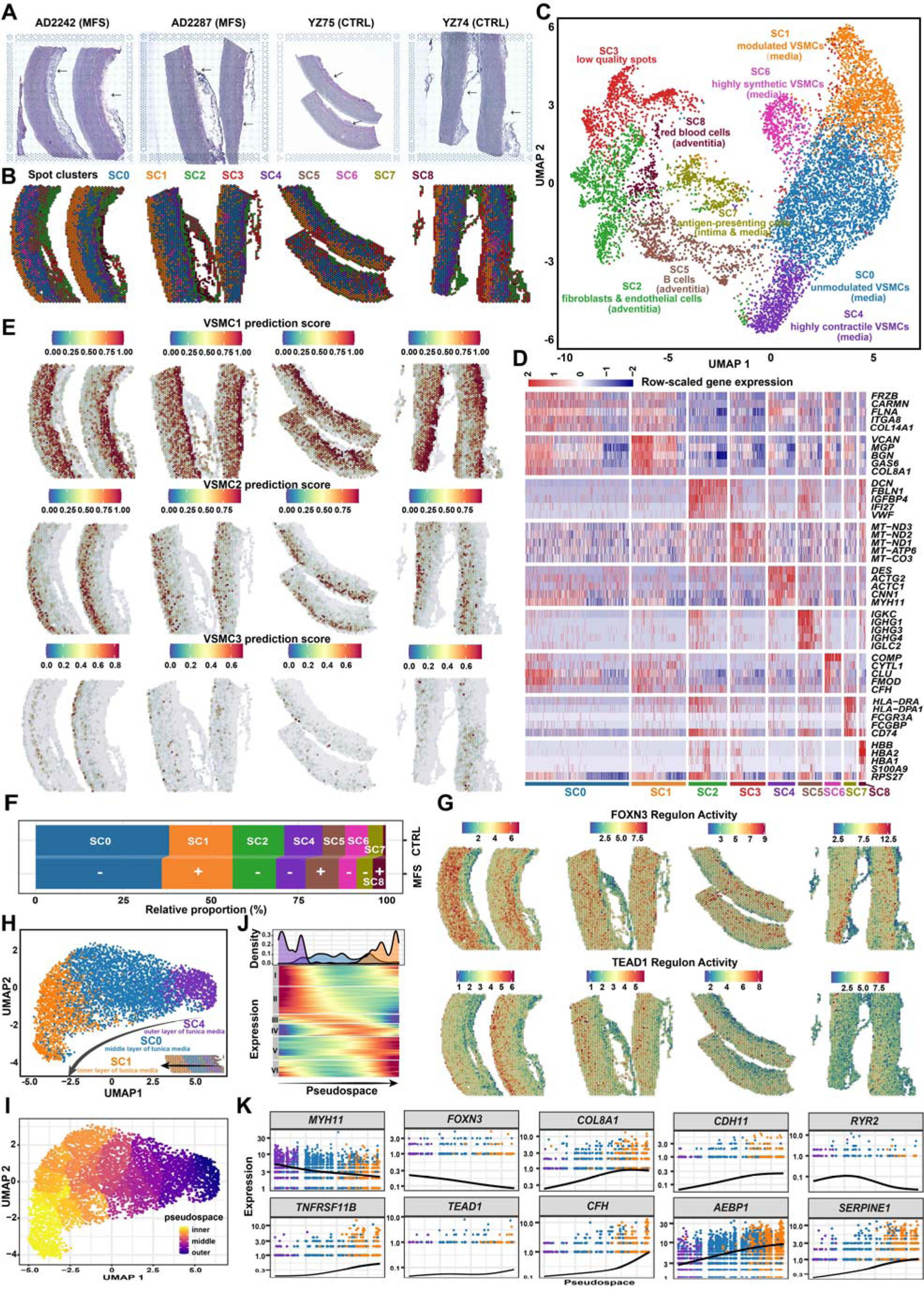
Spatially resolved transcriptome showing the phenotypic spectrum of VSMCs across the tunica media of human aortic root tissue. A) H&E staining images of aortic root tissue sections from the MFS and CTRL groups. The arrows indicate the tunica adventitia. Two sections for each subject and two subjects for each group were subjected to spatial transcriptomic assays. B) Unsupervised clustering of the spatial spots identified nine spot clusters. C) UMAP plot showing the nine spot clusters. Each cluster was annotated according to its expression profile and spatial location. D) Heatmap showing the expression of molecular features for each spot cluster. E) Spatial locations of the VSMC subclusters VSMC1, VSMC2, and VSMC3 on the tissue sections inferred by integrating the single-nucleus data and spatial transcriptomic data. The label transfer workflow of Seurat was applied in the prediction. F) Relative proportion of each spot cluster in each group. The average proportions of each group (n=2) are shown. SC3, which represents low-quality spots, was excluded from this analysis. +: expansion, -: contraction. G) Spatial distribution of the expression activities of the FOXN3 and TEAD1 regulons. H) UMAP plot showing the three major spot clusters over the tunica media. I) Pseudospace ordering of the three major spot clusters over the tunica media. J) Heatmap showing the expression profile of the genes that were expressed as a function of pseudospace. The significance threshold was set to a q-value < 0.05. The genes that changed as a function of pseudotime were detected with graph-autocorrelation analysis by using the “graph_test” function in Monocle3. K) Expression dynamics of markers and candidate regulators of VSMCs across the pseudospace.

### 2.7. FOXN3 may function as a key regulator for maintaining the contractile phenotype of human aortic VSMCs through targeting the *ACTA2* promoter

We proceeded with *in vitro* characterization of the functional role of the candidate key regulator FOXN3 in human aortic smooth muscle cells (HASMCs). PDGF-BB and TGFβ treatments are commonly utilized to induce dedifferentiation and enhance the contractile phenotype in VSMCs *in vitro*, respectively^[18]^. Of interest, PDGF-BB stimulation significantly decreased whereas TGFβ stimulation markedly increased both the mRNA and protein expression of FOXN3 and contractile markers including ACTA2 (α-SMA), TAGLN (SM22α), and CNN1 (calponin-1; Figure 6A,B; Figure S13, Supporting Information). This suggested that the expression of FOXN3 was positively correlated with the contractility of HASMCs. Next, siRNA-mediated knockdown of *FOXN3* was performed. The protein and mRNA expression of contractile markers was significantly decreased (Figure 6C; Figure S14A, Supporting Information) and the contractility of HASMCs was significantly reduced, as evidenced by the collagen gel contraction assay (Figure 6D). In addition, FOXN3 knockdown significantly increased the percentages of polygonal-shaped cells, i.e., modulated VSMCs (Figure 6E) and Ki-67-positive cells, i.e., proliferating cells (Figure 6F), indicating that the knockdown of FOXN3 promoted phenotypic modulation of HASMCs toward a dedifferentiated phenotype. Moreover, the analysis of bulk RNA-seq data revealed that the genes downregulated after *FOXN3* knockdown were notably enriched in GO terms related to muscle contraction, represented by *ACTA2*, *CNN1*, and *TAGLN*, and smooth muscle cell-matrix adhesion, represented by *DDR1*, *VTN*, and *PLA* (Figure S15; Table S16, Supporting Information). When treated with TGF-β, the knockdown of FOXN3 resulted in a significant increase in the proportion of cells with a polygonal shape, indicating an enhanced phenotypic modulation (Figure S16, Supporting Information). Additionally, while TGF-β promoted the proliferation of HASMCs, FOXN3 knockdown did not alter the effects of TGF-β on proliferation (Figure S16, Supporting Information).

**Figure 6.**
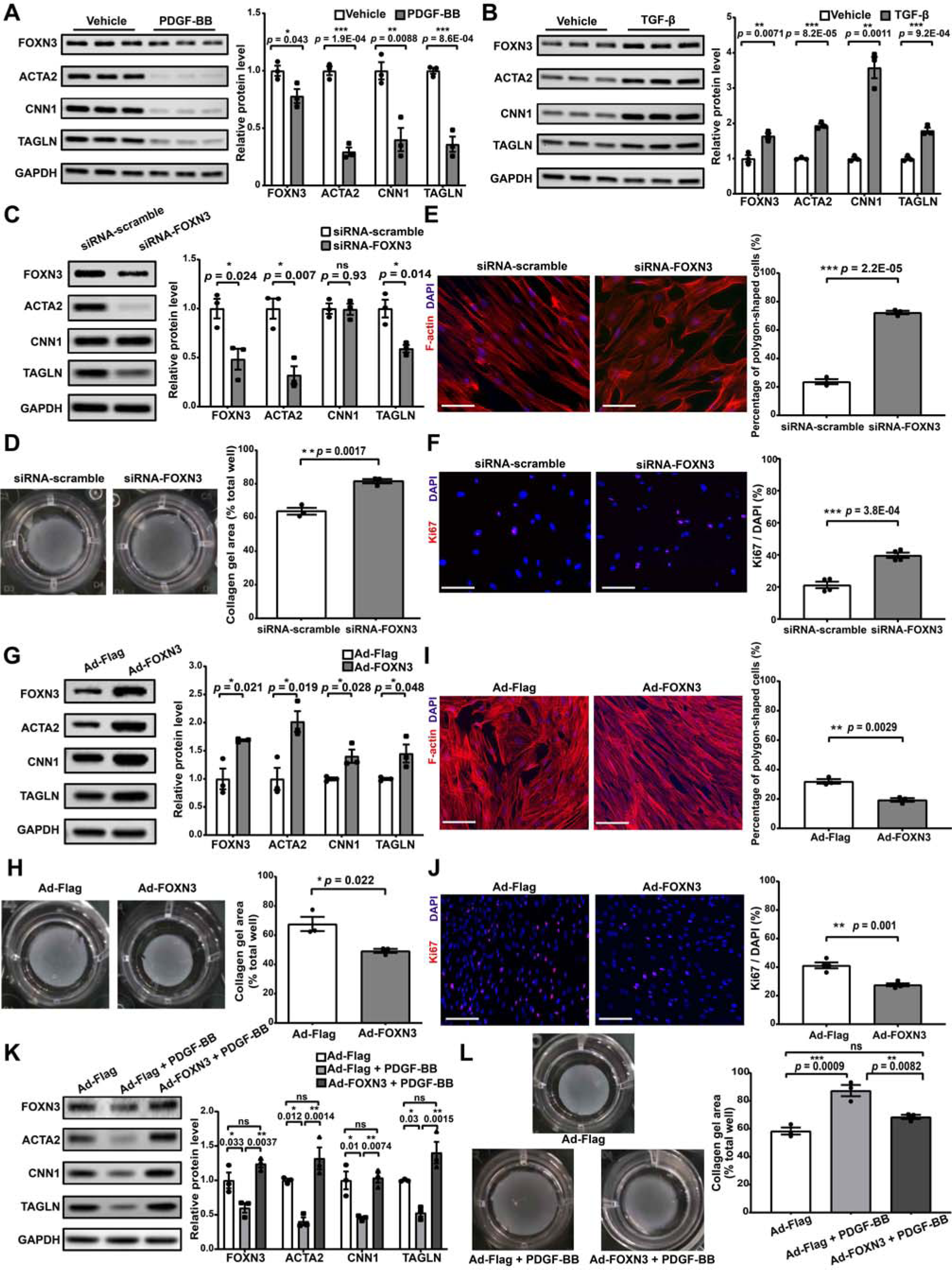
FOXN3 may function as a key regulator for maintaining the contractile phenotype of human aortic VSMCs. A) Western blot assay of FOXN3 and VSMC contractile marker proteins (ACTA2, CNN1, and TAGLN) in HASMCs following PDGF-BB treatment (20 ng/mL, 48 h post-treatment). B) Western blot assay of FOXN3 and VSMC contractile marker proteins in HASMCs following TGF-β treatment (10 ng/mL, 48 h post-treatment). C) Western blot assay of FOXN3 and VSMC contractile marker proteins in HASMCs transfected with scrambled siRNA (10 nmol/L) or FOXN3-siRNAs (10 nmol/L, 96 h post-transfection). D) Collagen gel contraction assay of HASMCs transfected with scrambled siRNA or FOXN3-siRNA (72 h post-transfection). E) Representative immunofluorescence staining images of F-actin (red) in HASMCs transfected with scrambled siRNA or FOXN3-siRNA (72 h post-transfection). F) Representative immunofluorescence staining images of Ki-67 (red) in HASMCs transfected with scrambled siRNA or FOXN3 siRNA (72 h post-transfection). G) Western blot assay of FOXN3 and VSMC contractile marker proteins in HASMCs infected with Adenovirus-FLAG-vector (Ad-Flag) or Adenovirus-FLAG-FOXN3 (Ad-FOXN3; 96 h post-infection). H) Collagen gel contraction assay of HASMCs infected with Ad-Flag or Ad-FOXN3 (96 h post-infection). I) Representative immunofluorescence staining images of F-actin (red) in HASMCs infected with Ad-Flag or Ad-FOXN3 (96 h post-infection). J) Representative immunofluorescence staining images of Ki-67 (red) in HASMCs infected with Ad-Flag or Ad-FOXN3 (96 h post-infection). K) Western blot assay of FOXN3 and VSMC contractile marker proteins in HASMCs infected with Ad-Flag or Ad-FOXN3 for 72 h and then subjected to PDGF-BB (20 ng/mL) treatment for 24 h. L) Collagen gel contraction assay of HASMCs infected with Ad-Flag or Ad-FOXN3 and then treated with PDGF-BB. In A-L, the data are presented as the mean ± SEM (three independent experiments). *: *p*-value < 0.05, **: *p*-value < 0.01, ***: *p*-value < 0.001, ns: not significant. The two-tailed Student’s t-test was used to compare two groups of data, while one-way ANOVA followed by multiple comparisons using Tukey’s method was used to compare multiple groups of data. In D, F, H, and J, the percentage of polygonal-shaped cells or Ki-67-positive cells in each image was calculated as the mean of the measurements in at least five representative views. Nuclei stained by DAPI are indicated in blue. Scale bar: 100 μm.

We also performed adenoviral-mediated overexpression of *FOXN3* and found that the protein and mRNA expression of contractile markers was significantly increased (Figure 6G; Figure S14B, Supporting Information) and that the contractility was significantly enhanced (Figure 6H). The percentages of polygonal-shaped cells and Ki-67-positive cells were significantly decreased (Figure 6I,J), suggesting that the overexpression of FOXN3 enhanced the contractile phenotype of HASMCs. Moreover, experimental evidence showed that the phenotypic modulation of HASMCs induced by PDGF-BB stimulation was attenuated by FOXN3 overexpression (Figure 6K,L; Figure S17, Supporting Information).

It is established that ACTA2 (also known as alpha-smooth muscle actin, α-SMA) is not only a widely utilized marker for contractile VSMCs, but also a key component of the contractile apparatus in VSMCs^[19]^. Mutations in ACTA2 cause familial thoracic aortic aneurysms and dissections^[20]^. Our single-nucleus data supported that ACTA2 may be a target of FOXN3 (Figure 4J). To validate this, CUT&Tag-qPCR was performed using the FOXN3 antibody in HASMCs (Figure 7A). The FOXN3 antibody significantly enriched the *ACTA2* promoter fragment, whereas IgG, used as a negative control, did not exhibit specific amplification. Moreover, the relative fold enrichment of the *ACTA2* promoter fragment further increased upon FOXN3 overexpression. These results demonstrated the binding of FOXN3 to the *ACTA2* promoter in HASMCs. In addition, luciferase reporter assay showed that FOXN3 increased the activity of the *ACTA2* promoter driving a luciferase reporter in HEK293A cells in a concentration-dependent manner (Figure 7B,C), suggesting that FOXN3 had regulatory effects on the activity of the *ACTA2* promoter.

**Figure 7.**
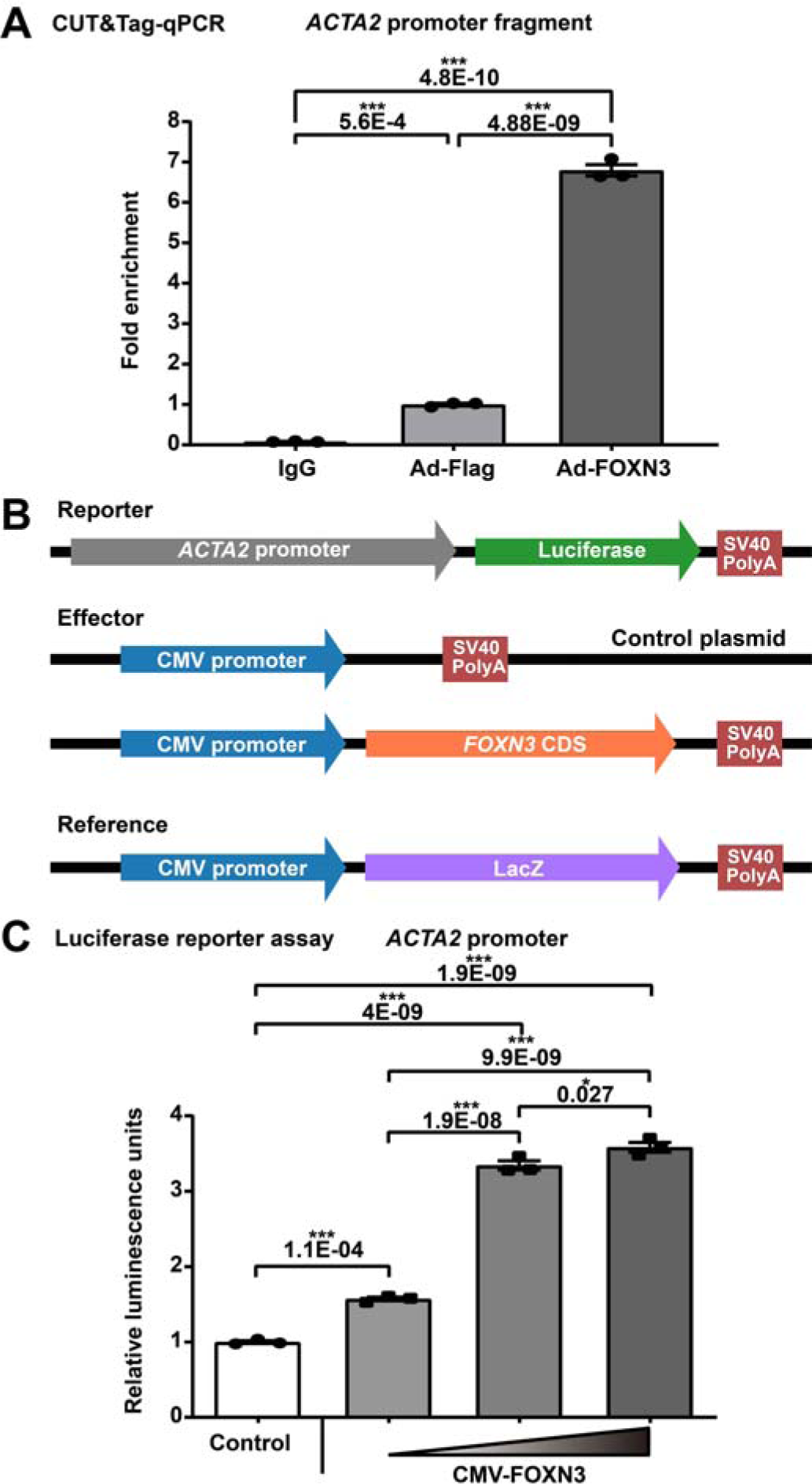
FOXN3 regulates smooth muscle contraction through targeting *ACTA2* that encodes the key component of the contractile apparatus in smooth muscle cells. A) CUT&Tag-qPCR experiment demonstrated the binding of FOXN3 to the *ACTA2* promoter region in HASMCs. IgG was used as a negative control. Ad-Flag: HASMCs infected with Adenovirus-FLAG-vector. Ad-FOXN3: HASMCs infected with Adenovirus-FLAG-FOXN3. B) Schematic diagram of reporter, effector, and reference plasmid construction for the luciferase reporter assay. C) FOXN3 increased the activity of *ACTA2* promoter driving a luciferase reporter in HEK293A cells in a concentration-dependent manner. In A and C, data are presented as mean ± SEM (n = 3 wells per group). *: *p*-value < 0.05, ***: *p*-value < 0.001. One-way ANOVA with Tukey’s multiple comparison correction.

Together, these results suggest that FOXN3 may function as a key regulator for maintaining the contractile phenotype of human aortic VSMCs through targeting the *ACTA2* promoter and thus may serve as a potential target for treating aortic aneurysms.

## 3. Discussion

Despite the performance of extensive research using animal models to gain insights into the disease progression, extrapolating animal data to humans has remained challenging^[21]^. Understanding the cellular and molecular dynamics of pathophysiological processes in human patient-derived specimens is of fundamental importance to the development of effective targeted therapy. Nevertheless, the complex cellular and regulatory landscape of tissues *in vivo* cannot be delineated with traditional bulk-level methods. In the present study, leveraging the state-of-the-art techniques including joint profiling of gene expression and chromatin accessibility in a single nucleus as well as spatial transcriptomics, we produced the first single-nucleus atlas of gene expression and chromatin accessibility in human aortic root tissues under healthy and MFS-associated aneurysmal conditions. Cell-type-specific transcriptomic and cis-regulatory profiles in the human aortic root were revealed. Cell-type-resolved regulatory changes in MFS patients compared with healthy controls were identified, particularly in the cardinal cell type, VSMCs. The gene expression and chromatin accessibility dynamics during the phenotypic modulation of VSMCs were uncovered. Moreover, candidate key regulators driving the phenotypic modulation of VSMCs were prioritized (such as *FOXN3*, *TEAD1*, *BACH2*, and *BACH1*). Finally, we showed experimental evidence supporting that FOXN3 functions as a novel key regulator for maintaining the contractile phenotype of human aortic VSMCs.

The discrepancies in embryological origin and hemodynamic features create a high degree of regulatory heterogeneity among different segments of the aorta including the aortic root, aortic arch, ascending thoracic aorta, descending thoracic aorta, and abdominal aorta^[22]^. In contrast to previous single-cell/nucleus transcriptome studies focusing on the human ascending aorta^[8,9]^, this study fills the knowledge gap caused by the lack of single-cell/nucleus datasets of the human aortic root in healthy and diseased conditions, although an scRNA-seq dataset of the aortic root tissue from only one MFS patient is publicly available (GEO accession: GSM4646673)^[7]^. In addition, to our knowledge, this study provides the first single-nucleus chromatin accessibility landscape in human aneurysmal aortic tissue (137,311 identified cCREs), although single-nucleus chromatin accessibility profiling has recently been applied to other human vascular tissues such as carotid arteries^[23]^ and coronary arteries^[12]^. Moreover, while most previous single-cell studies collected single-modality data, this study adopted simultaneous profiling of gene expression and chromatin accessibility in the same nucleus, allowing for robust definitions of cellular states and accurate reconstruction of the link between expression and accessibility^[24]^. Together, our study provided a unique and invaluable dataset that is expected to deepen our understanding of aortic biology and pathology and aid with, for example, the interpretation of risk variants identified by the ever-growing number of genome-wide association studies.

A growing body of evidence shows that multiple phenotypes of VSMCs exist even in healthy arteries of humans^[25]^. In this study, a considerable proportion (20∼30%) of modulated VSMCs with two distinct states (VSMC2 and VSMC1) were found to be present in the aortic root tissues of MFS patients and healthy controls (Figure 2B). This result reflects that phenotype modulation of VSMCs originates from the physiological response to maintain arterial wall homeostasis^[26]^. Although a statistically significant expansion of the modulated VSMCs was observed in MFS patients versus healthy controls, the cellular compositional alterations were generally modest (Figure 2A and Figure 3E). These results are quite similar to those observed in a single-nucleus transcriptomic study comparing ascending aortic tissue from patients with sporadic aortic aneurysms and healthy controls^[9]^. Therefore, these results represent the true cellular composition and alteration in humans. Despite MFS being regarded as an aggressive form of hereditary aortic aneurysm, the pathological stage at the time of prophylactic surgery and specimen collection is not late, which may also explain the modest compositional changes. Moreover, our data showed that the modulated VSMCs are not only associated with the diseased states (i.e., with an increased relative proportion in MFS) but also associated with the spatial location. The modulated VSMC subclusters are preferentially located in the inner layer of the tunica media close to the intima in both healthy and diseased conditions, as indicated by the spatial transcriptomic assay (Figure 5E). Compared to the outer-layer VSMCs, the inner-layer VSMCs may have to respond to stronger environmental stimuli and therefore tend to acquire a modulated synthetic phenotype. For example, inner-layer VSMCs can sense shear stress indirectly through endothelial cell-VSMC interaction^[26,27]^.

Leveraging the single-nucleus multiomic datasets, we, for the first time, disentangled the complex dynamics of gene expression and chromatin accessibility during the phenotypic modulation of human aortic root VSMCs (Figure 3F,G). Furthermore, based on the evidence from multiple comparative analyses, such as scWGNCA, regulon expression activity analysis, TF bind motif activity analysis, TF footprinting analysis, and pseudospace trajectory analysis (Figures 4 and 5), we obtained 11 candidate key regulators that potentially drive the phenotypic modulation of VSMCs and the pathogenesis of aortic root aneurysms. Among them, some have been experimentally demonstrated to play critical roles in the phenotypic modulation of VSMCs, reflecting the reliability of our single-nucleus multiomic date-driven candidate prioritization. For example, TEAD1, a member of the transcriptional enhancer activator domain (TEAD) TF family, represses the expression of contractile genes in VSMCs by abolishing the function of myocardin^[16]^. BACH1 (BTB and CNC homology 1), a member of the BACH TF family, represses the expression of contractile genes in VSMCs by suppressing chromatin accessibility at the promoters^[28]^. In contrast to TEAD1 and BACH1, which function as repressors of contractile gene expression, we found that the forkhead transcription factor FOXN3 may function as a key regulator for maintaining the contractile phenotype of human aortic VSMCs (Figure 6), whose expression decreased during phenotypic modulation and was downregulated in MFS versus CTRL VSMCs (Figure 4E). The functional role of FOXN3 has been investigated primarily in liver glucose metabolism^[29]^ or tumorigenesis^[30]^. However, the role of FOXN3 in the phenotypic modulation of VSMCs and the pathogenesis of aortic aneurysms has not been reported before. To our knowledge, this study represents the first report that FOXN3 functions as a regulator for maintaining the contractile phenotype of human aortic VSMCs probably through targeting the promoter of *ACTA2*. Nevertheless, the detailed regulatory mechanism of FOXN3 in VSMCs has yet to be elucidated.

This study is not without limitations. Due to the difficulty of obtaining specimens from healthy hearts, instead, we used the aortic root tissues of heart transplant recipients as controls. Although we recruited only heart transplant patients who suffered from heart failure caused by cardiomyopathy but had no aortic diseases, the extent to which heart failure affects the transcriptome of aortic root tissue remains to be determined. In addition, only MFS-associated aneurysmal conditions were considered in this study. The generality of the findings to other types of aortic root aneurysms, for example, sporadic forms, warrants further investigation. While no statistically significant difference was observed in the relative proportions of the subclusters for other cell types, except for VSMCs (Figure 2B), it is important to note that individual variations in cellular composition (Figure S8) and the relatively small sample size may diminish statistical power. A larger sample size in future studies would enhance the ability to detect potential compositional differences in other cell types, such as macrophages.

In summary, we presented the first atlas of gene expression and chromatin accessibility in the human aortic root at single-nucleus and spatial resolution. This unique dataset provides novel insights into the regulatory and spatial dynamics during phenotypic modulation in the aneurysmal aortic root of humans. FOXN3 was identified as a novel key regulator for maintaining the contractile phenotype of human aortic VSMCs, which may serve as a potential therapeutic target. Our datasets are expected to serve as a valuable reference to further decipher the regulatory mechanism of aortic root aneurysms and to interpret the risk loci for aortic root-related diseases in humans.

## 4. Experimental Section

### Ethics statement

All study procedures complied with the ethical regulations approved by the Ethics Committee of Fuwai Hospital, the Chinese Academy of Sciences (No. 2017-877). Written informed consent was provided by all enrolled subjects.

### Study subject enrollment

All patients enrolled in this study were diagnosed with MFS according to the Ghent II criteria and had undergone prophylactic aortic root replacement surgery at Fuwai Hospital. Genetic testing was conducted to exclude patients with features suggestive of other conditions, such as Loeys-Dietz syndrome. Patients with other aortopathy-related conditions, including aortitis, infection, aortic dissection, bicuspid aortic valve, and aortic atherosclerosis, were also excluded. Age- and ethnicity-matched heart transplant recipients without aortic diseases, but with heart failure caused by cardiomyopathy, were used as controls. Aortic root tissues were obtained during surgery from both groups for comparison.

### Tissue collection

For single-nucleus multiomic sequencing, aortic root tissues were collected during surgery, immediately frozen, and stored in liquid nitrogen until use for nuclei isolation. For spatial transcriptomic assays, fresh aortic root tissue was concurrently frozen in isopentane precooled by liquid nitrogen and embedded in optimum cutting temperature (OCT) compound.

### Single-nucleus multiomic sequencing

Nuclei were isolated and purified using an Shbio Nuclei Isolation Kit (52009-10, Shbio) according to the manufacturer’s instructions. Briefly, frozen aortic root tissue was thawed on ice, dissected into small pieces, and homogenized in LB solution with cold 1% BSA. After incubation on ice, the cell lysate was strained through a 40-μm filter and then spun down at 500 × g for 5 min at 4 [. Then, the supernatants were removed carefully, and the crude nuclei were resuspended in LB solution. PB1, PB2, and PB3 solutions were added sequentially to form 3 phases followed by centrifugation at 500 × g for 5 min at 4 [. The nuclei layer was aspirated and resuspended in NB twice and centrifuged at 500 g for 5 min at 4 [. The nuclei were sorted by staining and counted with Countstar Rigel S2. All buffers were supplemented with RNase Inhibitor (EO038, Thermo Fisher Scientific). Construction of single-nuclei ATAC and mRNA sequencing libraries was performed separately for each sample using a Chromium Next GEM Single Cell Multiome ATAC + Gene Expression Kit (v1, 10X Genomics) according to the manufacturer’s protocol. The libraries were sequenced using a NovaSeq 6000 (Illumina) for both mRNA and ATAC sequencing.

### Quality control of the single-nucleus multiomic sequencing data

Cell Ranger ARC (v2.0.0), the official analysis pipeline for 10X Chromium Single Cell Multiome ATAC + Gene Expression sequencing data, was used to perform read alignment (reference genome: refdata-cellranger-arc-GRCh38-2020-A-2.0.0), filtering, and counting. The gene-barcode expression matrix and ATAC fragment-barcode read count files were obtained for each sample. Subsequently, the R packages Seurat (v4.0.5) and Signac (v1.4.0) were used to jointly analyze the single-nucleus chromatin accessibility and gene expression data. To filter low-quality nuclei in each sample, we calculated quality metrics (including mRNA read count, ATAC read count, percentage of mitochondria gene reads, TSS enrichment score, and nucleosome signal) for each nucleus and filtered out the nuclei that did not meet the quality thresholds that we set for each sample (Table S2, Supporting Information). Scrublet (v0.2.3) was applied to further remove potential doublets.

### Peak calling and annotation

MACS2 (v2.2.7.1) was used to identify accessible sites (peaks) based on cell type-specific pseudobulk chromatin accessibility data. Peaks on nonstandard chromosomes and in genomic blacklist regions were removed using the functions “keepStandardChromosomes” and “subsetByOverlaps” of Signac, respectively. Each peak was annotated at multiple levels including the gene level, promoter level, exon/intron level, and exon level, with the R package ChIPpeakAnno (v3.34.1). The promoter was defined as the region 2,000 bp upstream and 500 bp downstream of the transcription start site (TSS). The “downstream” region of a gene was defined as the region 2,000 bp downstream of the gene body. The “upstream” region of a gene was defined as the region 5,000 bp upstream of the gene body. The cCREs were linked to genes by computing the correlation between their accessibility and the expression of nearby genes with the function “LinkPeaks” under default settings.

### Normalization, integration, dimensional reduction, and clustering of the single-nucleus multiomic sequencing data

For the gene expression data, each sample was individually normalized using the SCTransform procedure implemented in Seurat, and confounding sources of variation, including the percentage of mitochondria gene reads and cell cycle scores were regressed out. To correct for potential batch effects, gene expression data of all samples were integrated with the function “IntegrateData”. Then, the integrated data were subjected to dimensional reduction using principal component analysis (PCA).

For the chromatin accessibility data, following the selection of the most variable features (peaks) using the function “FindTopFeatures”, each sample was normalized using the term frequency-inverse document frequency (TF-IDF) normalization procedure followed by singular value decomposition (SVD). The above steps generated latent semantic indexing (LSI) embeddings, which were then used to integrate all datasets using the function “IntegrateEmbeddings” and to obtain new LSI embeddings corrected for batch effects.

To obtain a joint UMAP visualization that represents the measurements of both modalities, we computed a joint neighbor graph using the weighted nearest neighbor (WNN) methods implemented in the function “FindMultiModalNeighbors” of Signac. During this step, the top 30 PCA components of the gene expression data and the top 2-30 integrated LSI components of the chromatin accessibility data were taken as input. Unsupervised clustering of all nuclei was performed on the integrated gene expression data.

### Identification of cell-type/subcluster-specific cCREs or expressed genes

Cell-type/subcluster-specific cCREs or expressed genes were identified from the single-nucleus chromatin accessibility and gene expression data, respectively, using the function “FindAllMarkers” of the Seurat package (a specific cell type/subcluster versus all others). The test method used for differential accessibility was the logistic regression test (test.use = “LR”). The significance threshold for differential accessibility was set to a log2(fold change) value > 0.25 and a *p*-value adjusted for multiple testing < 0.01. The test method used for gene expression was the likelihood-ratio test (test.use = “bimod”). The significance threshold for gene expression was set to a log2(fold change) value > 0.5 and a *p*-value adjusted for multiple testing < 0.05.

### Differential gene expression analysis for the single-nucleus expression data

The DEGs between conditions in a specific cell type were identified using a method implemented in the R package Desingle (v1.20.0)^[31]^, which employed a zero-inflated negative binomial model to estimate the fraction of dropout and real zeros in the single-cell dataset. The following criteria were applied to consider a gene to be differentially expressed: absolute log2(fold change) value >1, adjusted *p*-value < 0.05, and categorization as “general differential expression” (significantly different expression between conditions concerning the expression abundance and the fraction of real zeros).

### Pseudobulk RNA-seq analysis

The raw UMI count matrix of the single-nucleus expression data in a specific cell type was summed per gene for each sample into a pseudobulk RNA-seq dataset. Differential expression analysis of the pseudobulk RNA-seq dataset was performed using the R package DESeq2 (v1.40.1) under the default settings. The statistical significance threshold was set to a *p*-value adjusted for multiple testing < 0.05.

### Differential accessibility analysis between conditions

To find differentially accessible cCREs between conditions, the function “FindMarkers” in the Seurat package was applied (test.use = ‘LR’, min.pct = 0.05, adjusted *p*-value < 0.05, logfc.threshold = 0.1).

### Functional enrichment analysis for a set of cCREs

To obtain functional interpretations for a set of cCREs, the Genomic Regions Enrichment of Annotations Tool (GREAT, v4.0.4)^[15]^ was applied under default settings, which analyzes the annotation of the nearby genes of the input cCREs. The significance threshold was set to a Bonferroni-corrected *p*-value < 0.05 (hypergeometric test).

### Functional enrichment analysis for a set of genes

Functional enrichment analysis for a set of genes was performed using CluoGO (v2.5.9)^[32]^ under default settings. The significance threshold was set to a Bonferroni-corrected *p*-value < 0.05.

### Gene set enrichment analysis

To perform GSEA for a cell type between conditions, we first ranked all the expressed genes by Signal2Noise (the difference in mean expression between the MFS and CTRL groups scaled by the standard deviation). Then, as input, the preranked gene list was imported to the GSEA software (v4.2.3). The precompiled REACTOME pathways in MsigDB (version: 7.5) were loaded. The significance cutoff was set to an FDR < 0.05. The EnrichmentMap (v3.4.4) plugin of Cytoscape (v3.9.1) was used to visualize the results with a network plot.

### TF binding motif enrichment analysis

To find TF motifs overrepresented in a given set of cCREs, we performed motif enrichment analysis using the hypergeometric test implemented in the function “FindMotifs” of the Signac package. The significance threshold was set to a *p*-value adjusted for multiple testing < 0.05.

### Differential TF motif activity analysis

A per-cell TF motif activity score was calculated using the R package chromVAR (v1.22.1), which detects motifs associated with variability in chromatin accessibility across cells. Then, the function “FindMarkers” of the Seurat package was used to identify TF motifs with significantly different activities between conditions (mean.fxn = rowMeans, fc.name = “avg_diff”, Wilcoxon rank-sum test, FDR < 0.05).

### TF footprinting analysis

For a given TF, the function “Footprint” of the Signac package was used to extract footprinting information for the TF motif from the chromatin accessibility data. The function “PlotFootprint” was used to visualize the footprinting by cell type or condition.

### Differential compositional testing

To detect statistically credible alterations in cellular composition derived from the single-cell dataset, we used a Bayesian approach implemented in scCODA (v0.1.9)^[33]^ (reference_cell_type = “automatic”, Hamiltonian Monte Carlo sampling method with default settings).

### Regulon expression activity analysis

Regulon activity analysis based on the single-nucleus expression data was performed following the tutorial of the R package SCENIC (v1.1.2)^[34]^. Briefly, gene coexpression modules in VSMCs were detected. Next, only the modules with significant enrichment of TF motifs were retained and referred to as regulons. Two databases “hg38 refseq-r80 10kb_up_and_down_tss.mc9nr.feather” and “hg38 refseq-r80 500bp_up_and_100bp_down_tss.mc9nr.feather” were utilized for the motif enrichment analysis. Finally, for each regulon, its activity was scored per nucleus. The activity scores of each regulon were compared between conditions, and the significance threshold was set to a Bonferroni-adjusted *p*-value < 0.05 (Wilcoxon rank-sum test, two-tailed).

### Pseudotime ordering of the single-nucleus expression data

To infer the trajectory of VSMC phenotypic modulation, we performed pseudotime ordering of the VSMC nuclei based on the single-nucleus expression data using the R package Monocle3 (v1.3.1) following the tutorial (https://cole-trapnell-lab.github.io/monocle3/). Then, the genes or cCREs that changed as a function of pseudotime were identified with graph-autocorrelation analysis (the “graph_test” function). The significance threshold was set to a q-value < 0.05 for gene expression and a q-value < 0.001 for chromatin accessibility.

### Single-cell weighted gene coexpression network analysis

To find functional gene modules of VSMCs, we performed scWGCNA based on the single-nucleus expression data using the R package scWGCNA (v0.0.0.9; https://github.com/Cferegrino/scWGCNA)^[35]^ following the tutorial. The module score was calculated per nucleus.

### Spatial transcriptomic assays

Sequencing libraries of spatial transcriptomic assays were prepared using a Visium Spatial Gene Expression Slide & Reagent kit (1000187, 10X Genomics) following the manufacturer’s instructions. Briefly, an OCT-embedded aortic root tissue section (10 μm) was placed on one of the capture areas (6.5 × 6.5 mm with ∼5000 barcoded spots) of a gene expression slide, and then stained with hematoxylin and eosin. A brightfield image was taken. After the tissue was permeabilized for the optimal time, reverse transcription was performed.

### Processing of the spatial transcriptomic data

The 10X Genomics official tool kit Space Ranger (v1.2.2) was utilized to perform sequencing read alignment, fiducial/tissue detection, and spot barcode/UMI counting of the spatial transcriptomic data for each section separately. The output gene-spot matrices of all sections were imported into Seurat (v4.0.5) for downstream analysis. The data were normalized for each section using the SCTransform procedure. Then, the data from different sections were integrated using the canonical correlation analysis procedure to correct for technical differences. After linear dimensional reduction was performed using principal component analysis (PCA), an SNN graph was constructed with the first 20 PCA components. The clustering of spots was performed using the Louvain algorithm (resolution: 0.4). UMAP dimensional reduction was conducted to visualize the spatial spots. To integrate the spatial transcriptomic data with the single-nucleus data, the label transfer workflow of Seurat was applied to assign each spot a prediction score for each subcluster obtained from the single-nucleus data analysis. The expression activity of a given gene set/pathway in each spot was quantified by calculating an activity score using the method implemented in Single Cell Signature Explorer (v3.3)^[36]^. To unravel the spatial transcriptomic dynamics across the tunica media of the aortic root tissue, we performed pseudospace ordering of the tunica media spots (spot clusters sc0, sc1, and sc4) using Monocle3. Genes whose expression changed as a function of the pseudospace were identified (the “graph_test” function) with a significance threshold q-value < 0.05.

### Cell culture and treatment

HASMCs were purchased from Lonza (CC-2571, Lonza). The HASMCs were maintained in SmBM^TM^ Basal Medium (CC-3181, Lonza) supplemented with SmGMTM-2 SingleQuots^TM^ supplements (CC-4149, Lonza) in a humidified 5% CO_2_ incubator at 37 °C according to the manufacturer’s instructions. Following serum starvation for 24 h, HASMCs were treated with TGF-β (10 ng/mL, 100-21, PeproTech) or PDGF-BB (20 ng/mL, 100-14B, PeproTech) for 24 h or 48 h, respectively.

### Quantitative real-time PCR

Total RNA was extracted from treated HASMCs using TRIzol reagent (15596018, Invitrogen). Gene expression was quantified with an SYBR Green real time-PCR using the ViiA™ 7 system (4453535, Thermo Fisher Scientific). β-Actin was used as a housekeeping gene to normalize the amounts of cDNA in each sample. The primer sequences were as follows: *FOXN3* (F: 5’-TCGTTGTGGTGCATAGACCC-3’, R: 5’-GTGGACCTGATGTGCTTTGATA-3’), *ATAC2* (F: 5’-CGTGCTGGACTCTGGAGATG-3’, R: 5’-GCCAGATCCAGACGCATGAT-3’), *TAGLN* (F: 5’-CCGTGGAGATCCCAACTGG-3’, R: 5’-CCATCTGAAGGCCAATGACAT-3’), *CNN1* (F: 5’-CTGTCAGCCGAGGTTAAGAAC-3’, R: 5’-CCATCTGAAGGCCAATGACAT-3’), and *ACTB* (F: 5’-GAGAAAATCTGGCACCACACC-3’, R: 5’-GGATAGCACAGCCTGGATAGCAA-3’).

### Western blot assay

The HASMCs or the tunica media of the aortic root tissue was homogenized in ice-cold RIPA buffer (P0013B, Beyotime) supplemented with protease and phosphatase inhibitor cocktail (P1046, Beyotime). The total protein concentration was quantified by BCA assay (23227, Thermo Fisher Scientific). SDS[PAGE was used to separate the proteins, which were subsequently transferred onto nitrocellulose membranes (66485, PALL). The membranes were blocked in 5% nonfat milk (P0216, Beyotime) for 1 h at room temperature. Then, the membranes were incubated overnight at 4 [with primary antibody diluted in 5% nonfat milk or primary antibody dilution buffer (P0256, Beyotime). Subsequently, the membranes were incubated for 1 h with an HRP-conjugated secondary antibody diluted at 1:10000 in 5% nonfat milk or secondary antibody dilution buffer (P0258, Beyotime) and detected with a BeyoECL Star Kit (P0018A, Beyotime). The following primary antibodies were used: GAPDH (1:2000, 2118L, Cell Signaling Technology), FOXN3 (1:1000, 711585, Thermo Fisher Scientific), ACTA2 (α-SMA, 1:1000, ab5694, Abcam), CNN1 (Calponin-1, 1:15000, ab46794, Abcam), β-Actin (1: 100000, 66009-1-Ig, Proteintech), and TAGLN (SM22α, 1:8000, ab14106, Abcam) antibodies. Western blot images were acquired using the Quantity One software (v4.6.9). Densitometry analysis was performed by quantifying the intensity of bands using the ImageJ software (v1.53e).

### Gene knockdown

HASMCs (passages 4–5) were seeded in 6-well plates the day before transfection. The cells were starved in Opti-MEM™ I Reduced-Serum Medium (31985062, Thermo Fisher Scientific) for 4 h before the transfection. Then, scrambled siRNA or FOXN3 siRNA (10 nM) was transfected into the cells using Lipofectamine RNAiMAX Reagent (13778150, Thermo Fisher Scientific) according to the manufacturer’s protocol. The sequences were as follows: siRNA-scramble (5’-UUCUCCGAACGUGUCACGU-3’) and siRNA-FOXN3 (5’-GGAGUCAGAGUAUUGGGAATT-3’). Transfection efficiency was evaluated via qRT[PCR at 48 h post-transfection and Western blot assay at 96 h post-transfection.

### Adenoviral-mediated gene overexpression

HASMCs (passages 4–5) were seeded in 6-well plates the day before infection. Cells were infected with adenovirus-FLAG-vector (Ad-Flag) or adenovirus-FLAG-FOXN3 (Ad-FOXN3) (HH20220630GX-AD01, HanBio) at a multiplicity of infection (MOI) of 50 for 96 h. For PDFG-BB treatment, the infected cells were challenged with PDGF-BB (20 ng/mL) for 48 h.

### Collagen gel contraction assay

Before the assay, 24-well culture plates were precooled on ice. HASMCs (5×10^5^ cells/ml) were suspended in DMEM/F12 (11320033, Thermo Fisher Scientific) supplemented with 2% FBS. Then, cell suspension (100 μL) was mixed with collagen (140 μL, 354236, Corning), culture medium (200 μL), and of 0.1 M NaOH solution (40 μL). After collagen polymerization, culture medium (500 μL) was added on top of each collagen gel lattice. After 24 h, the contractility of the gel was quantified by measuring the gel diameter.

### smFISH

Paraffin sections of aortic root tissue (10 μm) were baked and deparaffinized. Then, epitope retrieval, protease treatment, and probe hybridization were performed using highly sensitive RNAscope^®^ technology according to the manufacturer’s instructions. The RNAscope® Multiplex Fluorescent Reagent Kit v2 (323100) was applied to visualize hybridization signals. Images were acquired using the Vectra® Polaris™ software (v1.0.7). Then, the images were analyzed using the Phenochart software (v1.0.9). The RNAscope™ probes were as follows: Hs-MYH11 (444151-C3), Hs-RYR2 (415831), Hs-LAMA2 (530661), negative control (321831), and positive control (321811).

### Immunofluorescence staining

HASMCs were fixed in 4% paraformaldehyde and permeabilized with 0.5% Triton X-100 for 20 min. The cells were washed 3 times using 1×PBS and blocked with 3% BSA for 1 h at room temperature. Then, the cells were stained with antibodies against Ki67 (1:500, 9129S, Cell Signaling Technology) at 4 °C overnight followed by an Alexa Fluor 594-conjugated secondary antibody (1:500) (A-11012, Thermo Fisher Scientific). Immunofluorescence staining images were acquired using the Pannoramic Scanner software (v3.0.3.139795 RTM). The images were analyzed using the Slide Viewer software (v2.5) and the Image-Pro Plus software (v6.0.0.260).

### F-actin staining

Cells were fixed with freshly prepared 4% paraformaldehyde for 15 min and then permeabilized with 0.1% Triton X-100 in PBS for 15 min. Subsequently, the cells were incubated with rhodamine-phalloidin (1:400, R415, Thermo Fisher Scientific) for 60 min at room temperature. The nuclei were stained with DAPI (4083S, Cell Signaling Technology).

### H&E and EVG staining

The sections of formaldehyde-fixed paraffin-embedded aortic root tissues were baked and deparaffinized. To perform H&E staining, the sections were exposed to hematoxylin for 5 min, followed by differentiation using 1% hydrochloric acid alcohol. The sections were then placed in eosin staining solution for 1 min. Finally, the sections were dehydrated using graded ethanol, vitrified using dimethylbenzene, and mounted in synthetic resin. EVG staining (DC0059, Leagene Biotechnology) was performed according to the manufacturer’s instructions. Briefly, the sections were stained in Verhoeff working solution for 15 min, followed by differentiation using Ferric Chloride Differentiating Solution. The slides were then rinsed by two changes of 95% alcohol before being moved into Van Gieson’s Solution for 2-5 min. Finally, the sections were dehydrated in absolute alcohol and mounted in synthetic resin. The percentage of elastin area was calculated by the software ImageJ (v1.53e).

### Bulk RNA-seq

After incubating for 48 h, the cells were collected for RNA extraction using TRIzol reagent (15596018, Invitrogen). For the construction of bulk RNA-seq libraries, the NEBNext Ultra RNA Library Prep Kit for Illumina (E7530L, NEB) was utilized. The libraries underwent sequencing using the Illumina X Ten system. To ensure quality, fastp (v0.19.6) was employed for read quality control. Transcript abundance quantification was performed using kallisto (v0.45.0). Differential expression analysis between groups was carried out using the R package sleuth (v0.30.0). We defined the statistical significance threshold as a q value less than 0.05. Moreover, we established the biological significance threshold by requiring the absolute value of log2 (fold change) to be greater than 1.

### CUT&Tag-qPCR

Before the CUT&Tag assay, HASMCs at passages 4 were infected with adenovirus-FLAG-vector (Ad-Flag) or adenovirus-FLAG-FOXN3 (Ad-FOXN3) (HH20220630GX-AD01, HanBio) at an MOI of 50 for 96 h. Subsequently, 100,000 cells were processed using the Hyperactive Universal CUT&Tag assay Kit (TD904, Vazyme), following the manufacturer’s instructions. Primary antibodies against FOXN3 or IgG isotype control were incubated overnight. The secondary antibody was then incubated for 1 h at room temperature. Binding of the hyperactive pA/G-Transposon Pro with the secondary antibodies was carried out for 1 h. After fragmentation, DNA fragments were extracted, and qPCR was performed. The fold enrichment of DNA fragments was normalized with DNA spike-in. The following antibodies were used: FOXN3 (711585, Thermo Fisher Scientific; 0.5 µg for each reaction), rabbit IgG isotype control (ab37415, Abcam; 0.5 µg for each reaction). The primer sequences for the *ACTA2* promoter were as follows: Forward: 5’-AGCAAAAGGGGTTAAGGATGGG-3’, Reverse: 5’-GATGGGTGGGGAGCTGTTTT-3’. The primer was designed according to the prediction of FOXN3-binding sites in the *ACTA2* promoter region using JASPAR software.

### Luciferase Reporter Assay

The pGL4.11 Luciferase Reporter Vector (E6661, Promega) was used to clone the nucleotides −2000 to +99 relative to the transcription start site of the human *ACTA2* locus. HEK293A cells were cultured in DMEM supplemented with 10% FBS in 12-well plates. Cells were transfected with three plasmids, namely CMV-FOXN3, ACTA2-luc luciferase reporter, and β-galactosidase (β-gal) plasmid using Lipofectamine 3000 Reagent (L3000015, Thermo Fisher). A total of 1200 ng of plasmid DNA was consistently transfected. The ratios of CMV-FOXN3: ACTA2-luc luciferase reporter: β-gal plasmids were adjusted to 1:1:0.4, 2:1:0.4, or 3:1:0.4. Luminescence measurements were taken 48 h post-transfection using the Luciferase Reporter Assay System (DD1201, Vazyme) on the TECAN Infinite 200 PRO Plate reader. The relative firefly luciferase activity was normalized to β-gal activity.

### Statistical analysis

Statistical analyses were performed in R. Two-tailed student’s t-test or Wilcoxon rank-sum test was used to compare two groups of data. One-way analysis of variance (ANOVA) was used to analyze data among multiple groups. Following ANOVA, a post hoc test was performed using Tukey’s method. A *p*-value < 0.05 was considered statistically significant.

## Acknowledgments

We thank Qingzhi Wang at the Department of Pathology, Fuwai Hospital for technical support in cryosectioning and staining. This project was supported by grants from the National Natural Science Foundation of China (82170506), the National High Level Hospital Clinical Research Funding (2023-GSP-RC-21, 2022-GSP-GG-6, and 2022-PUMCH-C-025), the Open Projects of the State Key Laboratory for Cardiovascular Diseases (2022KF-04), and the Yunnan Provincial Cardiovascular Disease Clinical Medical Center Project (No. FZX2019-06-01).

## Author Contributions

X. L and Q. Z. contributed equally to this work. X. L. designed the project, analyzed the data, and wrote the manuscript. Q. Z. performed all the wet-lab experiments with the assistance of K. Y., Z. P., and K. W, and participated in drafting the manuscript. H. Y. and Q. C. contributed to specimen collection and molecular diagnosis. W. L. designed the web-based interface. M. L. and C. S. were responsible for subject enrollment and helped to interpret the results. M. L. and Z. Z. supervised the project and were responsible for the acquisition of funding.

## Data Availability Statement

All raw sequencing data have been deposited in Genome Sequence Archive for humans (https://ngdc.cncb.ac.cn/gsa-human/) and are available via accession number: HRA004063.

