## Supplementary material for "Single-nucleus Multiomic Analyses Identifies Gene Regulatory Dynamics of Phenotypic Modulation in Human Aneurysmal Aortic Root": Figure S1-S17

### SUPPLEMENTAL FIGURES

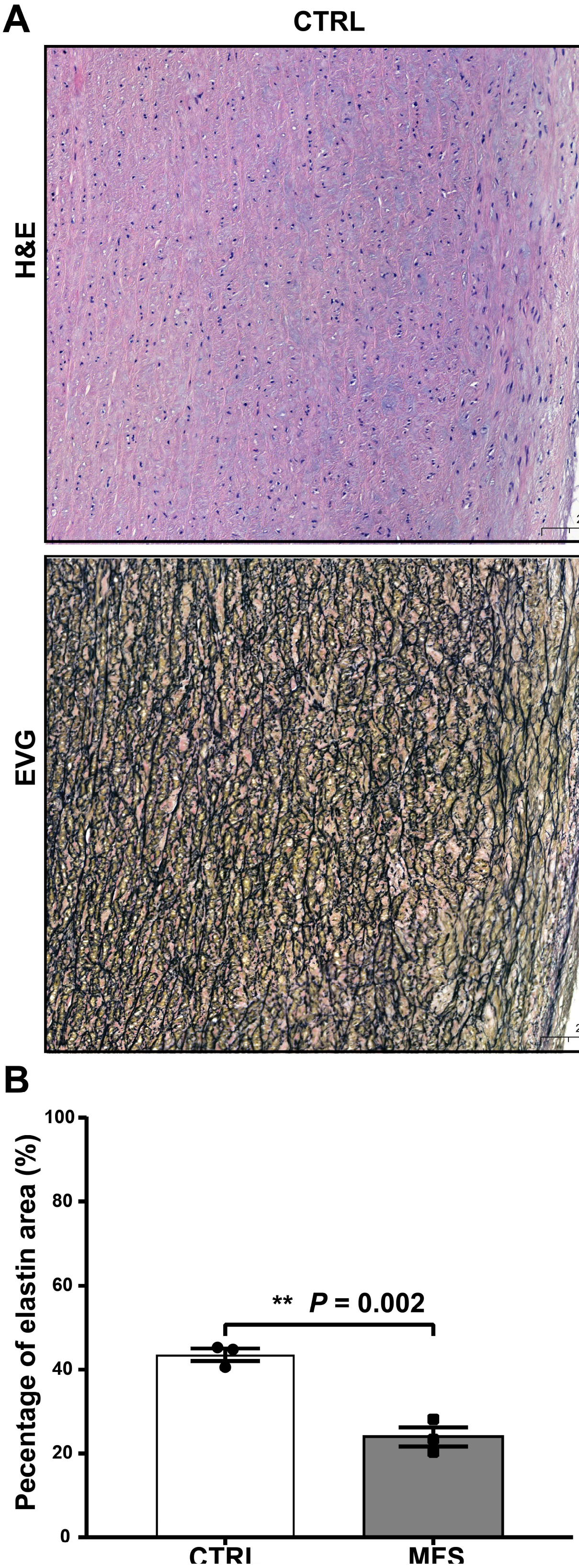


**Figure S1.** The H&E and EVG staining of aortic root tissue sections from MFS patients exhibited characteristic histopathological features of aortic aneurysms. A) Representative staining images. Consecutive sections from a single subject of each group are shown. Medial layer degeneration was observed in aortic root tissue sections from MFS patients, characterized by elastic fiber fragmentation or loss, mucoid extracellular matrix accumulation, and smooth muscle nuclei loss. The region highlighted by the red box shows notable signs of this degeneration. B) The percentage of elastin area stained with EVG was significantly decreased in individuals with MFS compared to the control group. **: $p$-value < 0.05, two-sided Student's t-test (n=3). MFS: Marfan syndrome; CTRL: control.

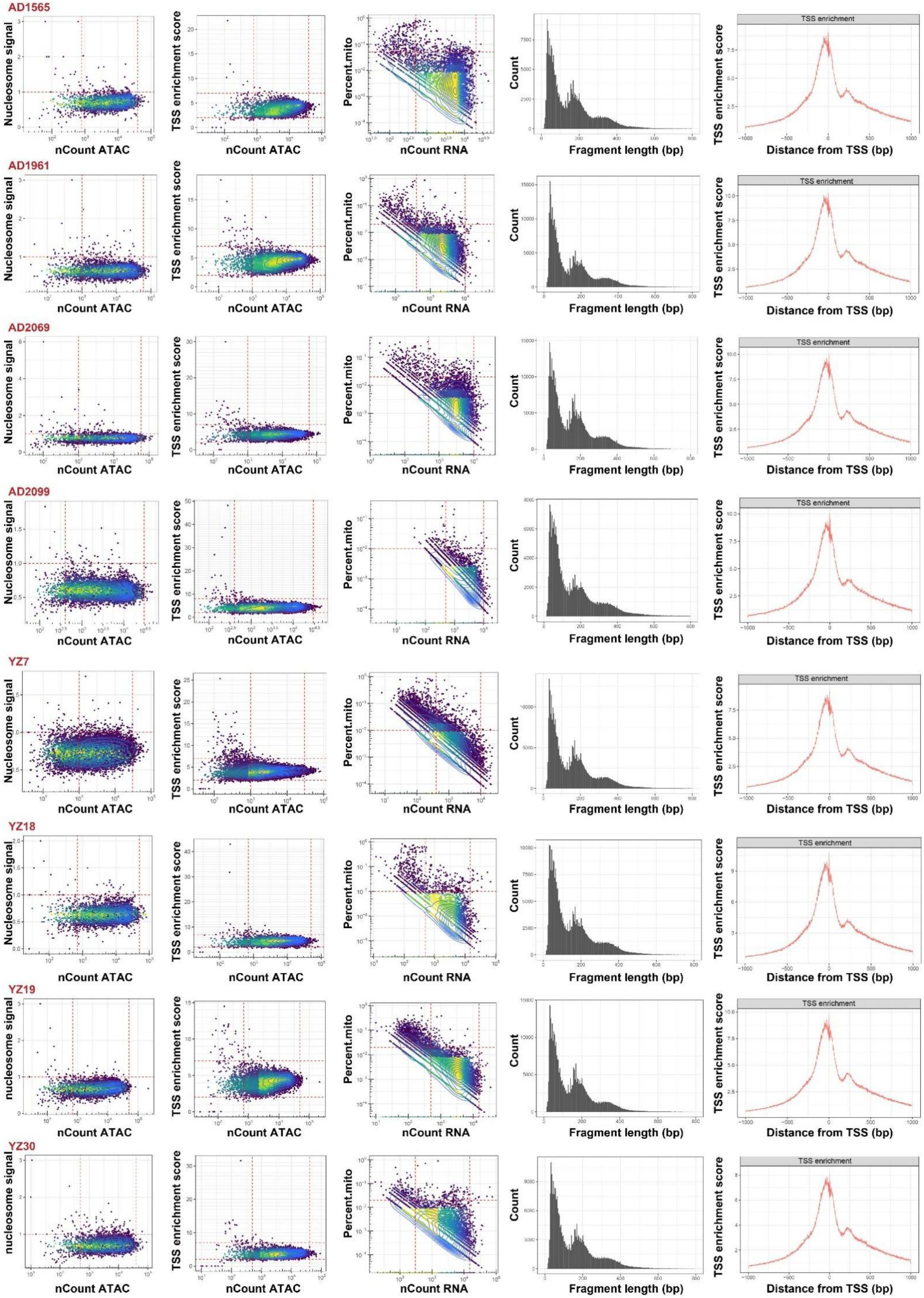

AD1565
AD1961
AD2069
AD2099
YZ7
YZ18
YZ19
YZ30
Nucleosome signal
nucleosome signal
nCount ATAC
TSS enrichment score
Percent.mito
nCount RNA
Count
Fragment length (bp)
TSS enrichment
Distance from TSS (bp)

**Figure S2.** Quality control of the single-nucleus multiomic sequencing dataset. The red dot line indicates the threshold used during the quality control step. TSS: transcription start site.

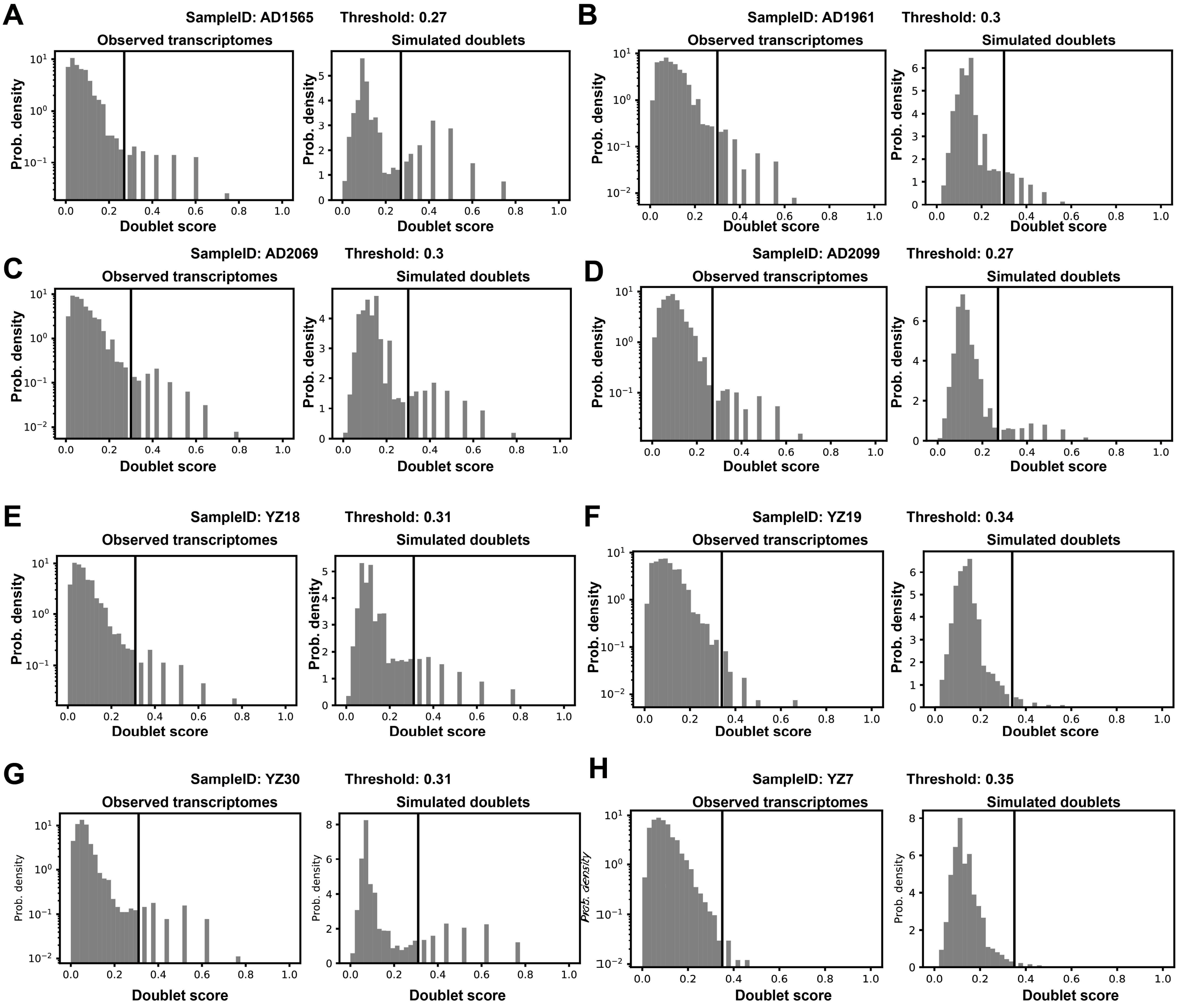


**Figure S3.** Histograms of doublet scores calculated by the python package Scrublet for observed transcriptomes (left panel) and simulated doublets (right panel) in each of the eight samples (A-H). The threshold for doublet prediction was determined based on the histograms and is indicated by the vertical line on the plot.

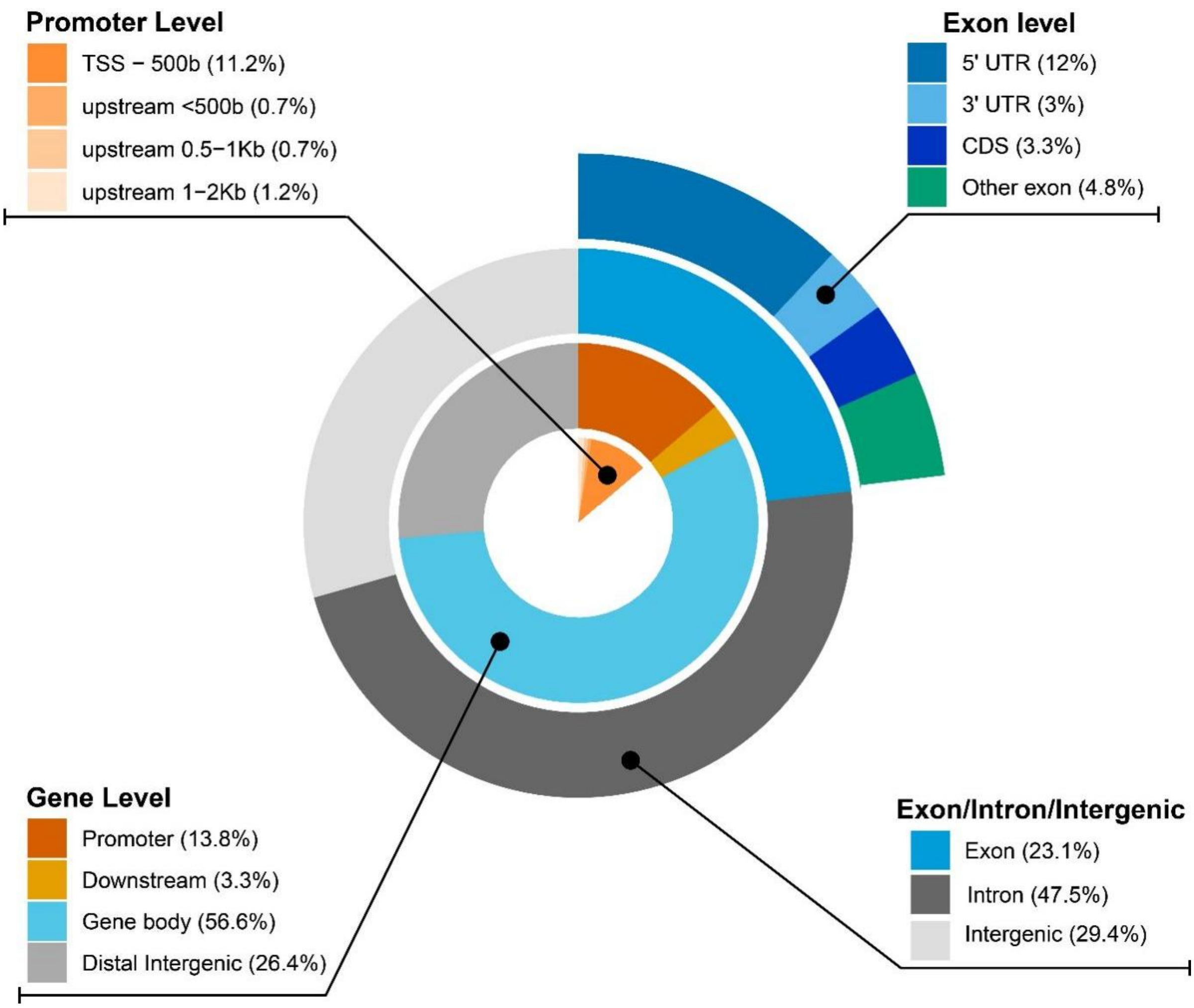


**Figure S4.** The proportion of each category for all cCREs. The annotation of cCREs was performed using the R package ChIPpeakAnno. The promoter was defined as the region 2,000 bp upstream and 500 bp downstream of the transcription start site (TSS). The “downstream” of a gene was defined as the region 2,000 bp downstream of the gene body. The “upstream” of a gene was defined as the region 5,000 bp upstream of the gene body.

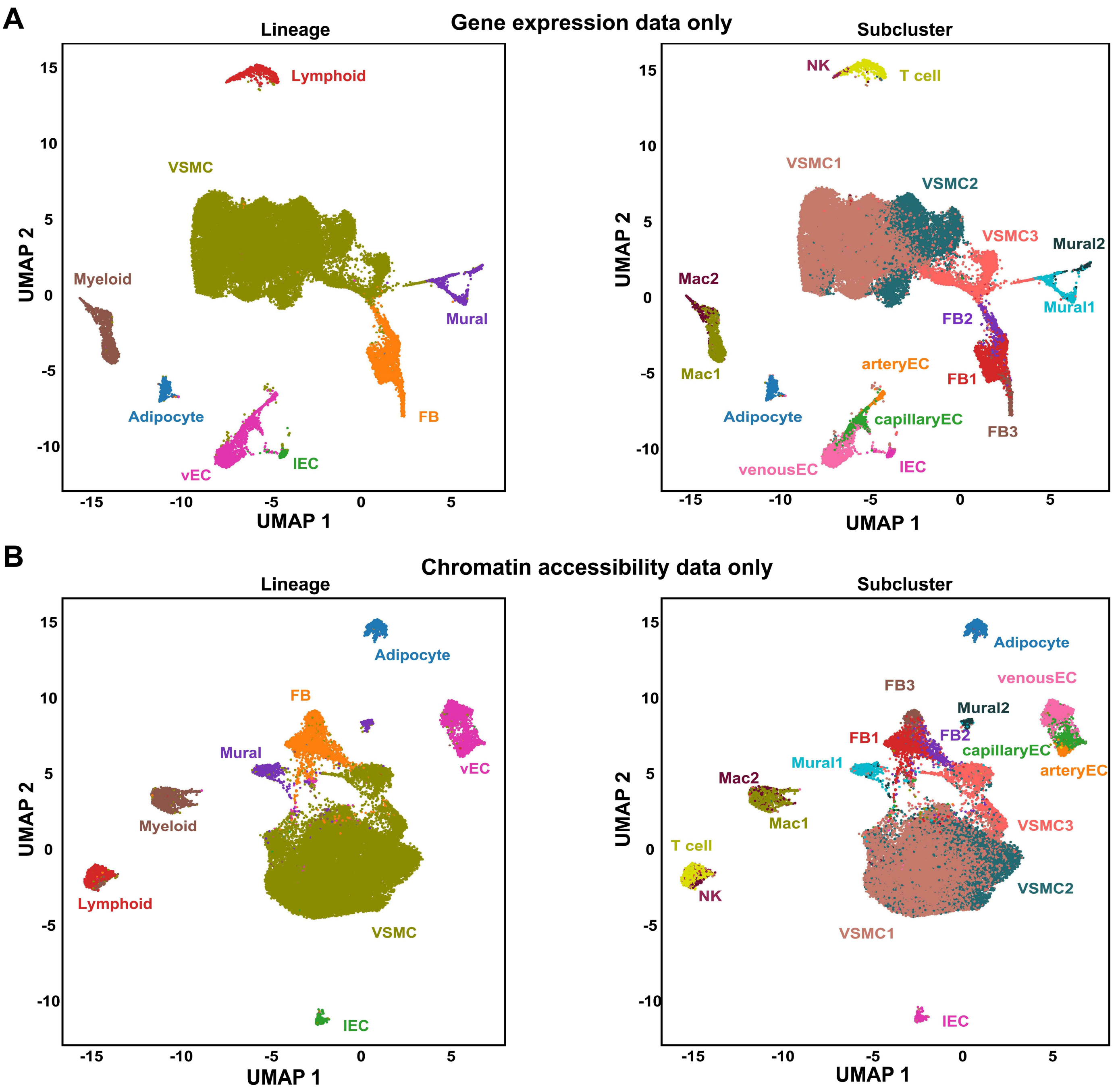


**Figure S5.** UMAP plots showing all nuclei projected in a 2D UMAP space based on only one modality of the datasets. A) UMAP plots showing all nuclei projected in a 2D UMAP space based on the gene expression data. B) UMAP plots showing all nuclei projected in a 2D UMAP space based on the chromatic accessibility data.

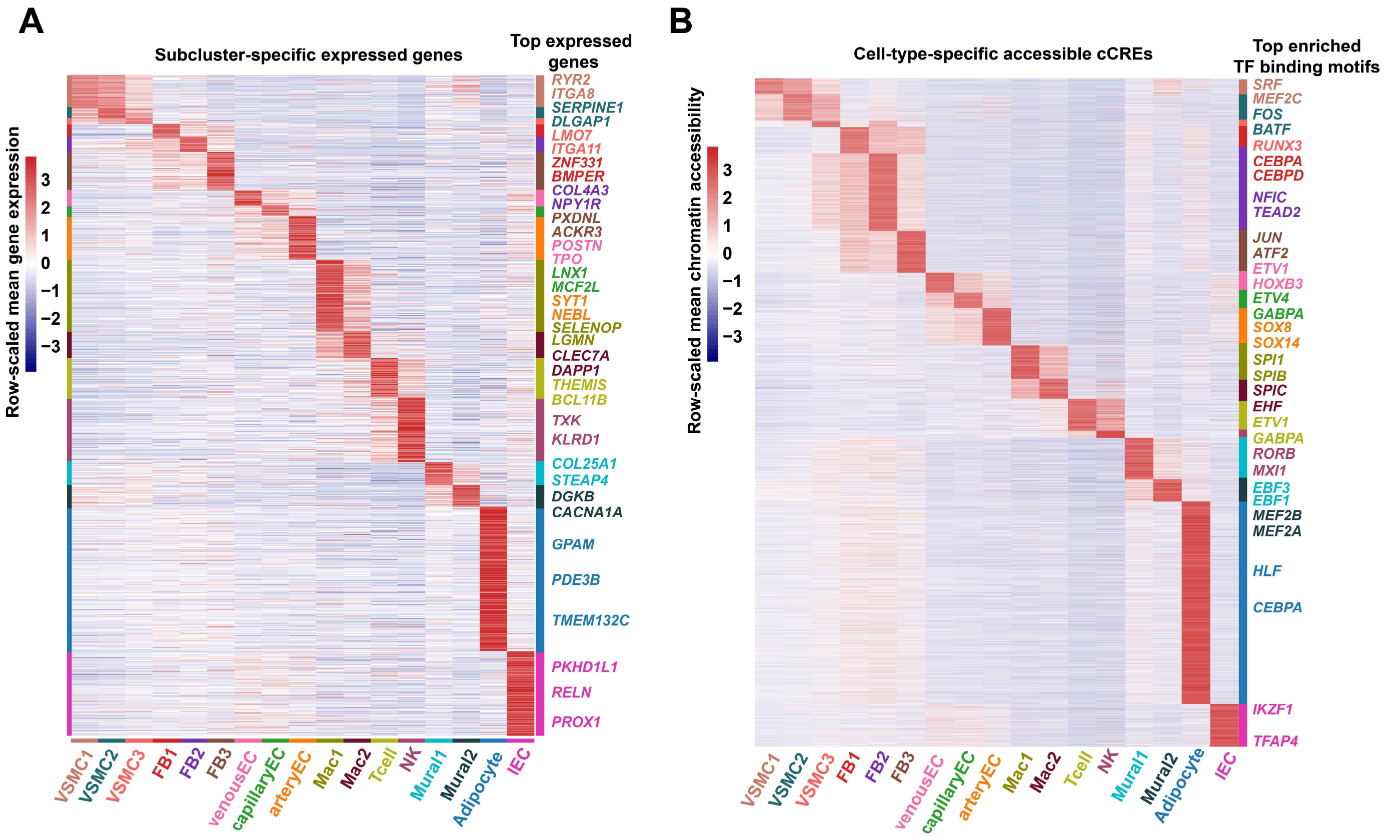


**Figure S6.** Heatmaps showing the profiles of subcluster-specific expressed genes (A) and assessable cCREs (B). The significance threshold for gene expression was set to a log2(fold change) value > 0.5 and a $p$-value adjusted for multiple testing < 0.05 (likelihood-ratio test). The significance threshold for differential accessibility was set to a log2(fold change) value > 0.25 and a $p$-value adjusted for multiple testing < 0.01 (logistic regression test). The cell-type-specific expressed genes or assessable cCREs were detected using the function “FindAllMarkers” of the Seurat package.

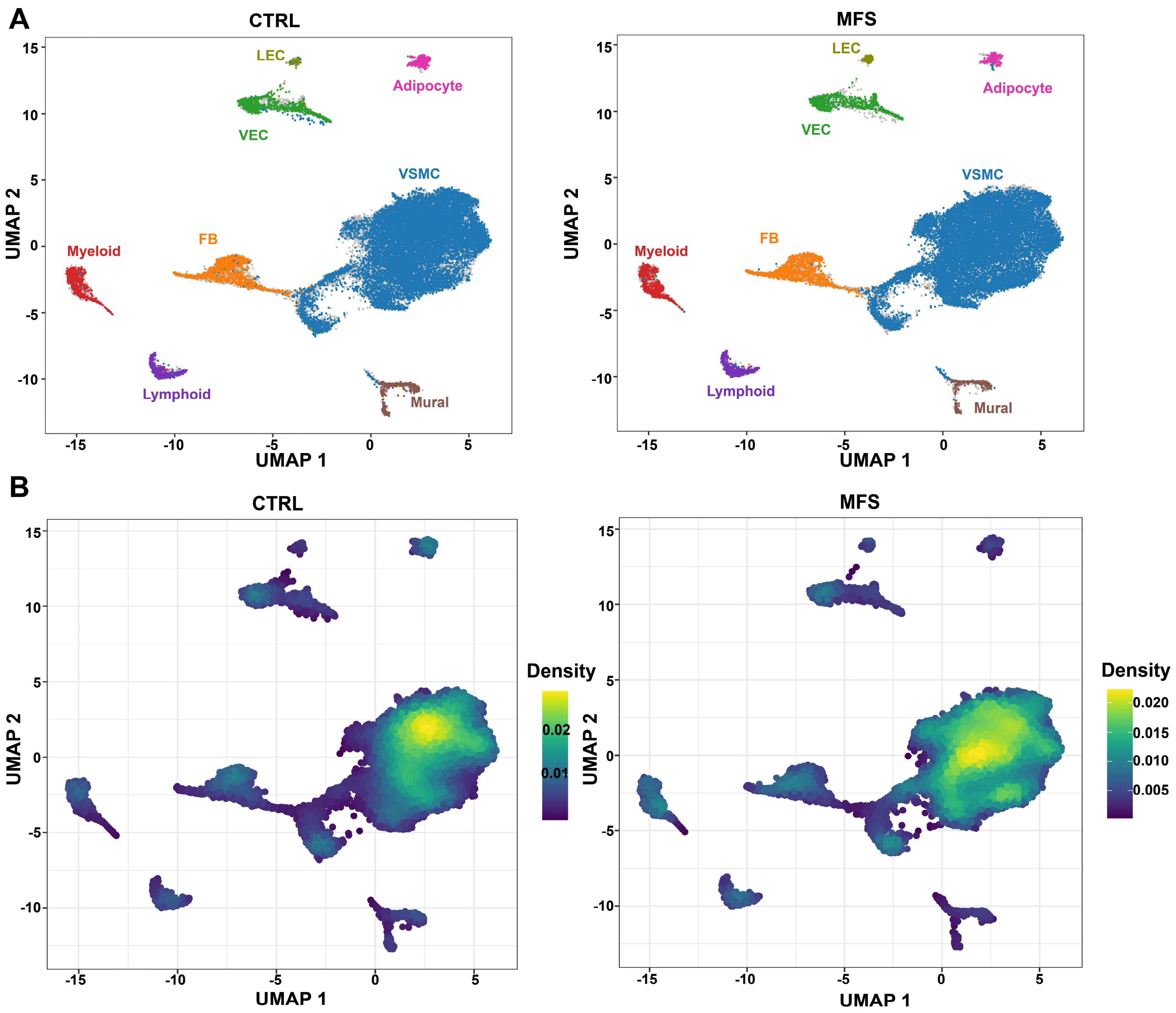


**Figure S7.** The nuclei from the CTRL or MFS group projected in a UMAP space. A) UMAP plot showing the distribution of nuclei from the CTRL group (left panel) or the MFS group (right panel). The nuclei are color-coded by cell type. B) UMAP plots showing the density of nuclei from the CTRL group (left panel) or the MFS group (right panel).

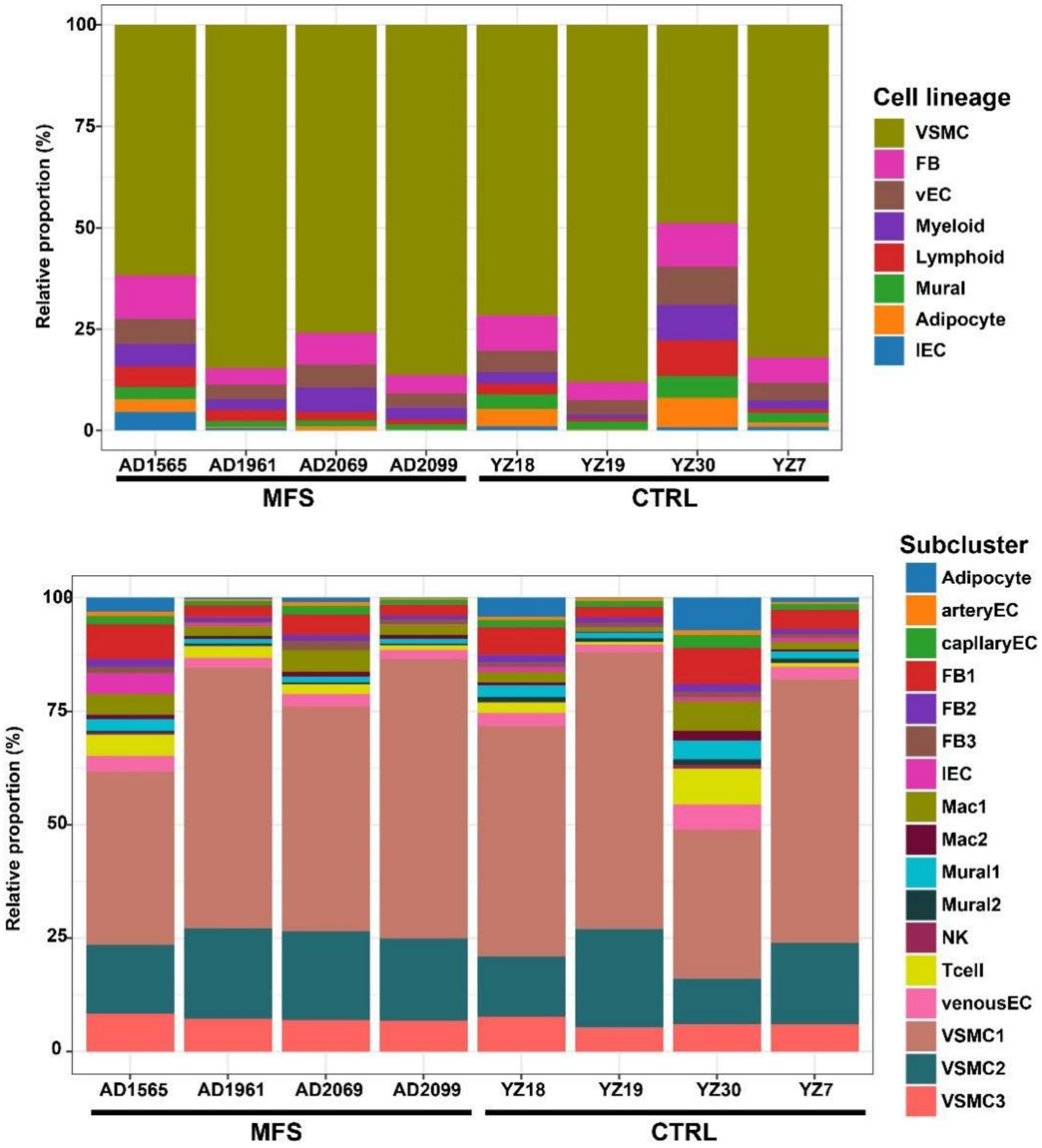


**Figure S8.** The relative proportion of each cell type/subcluster in each sample.

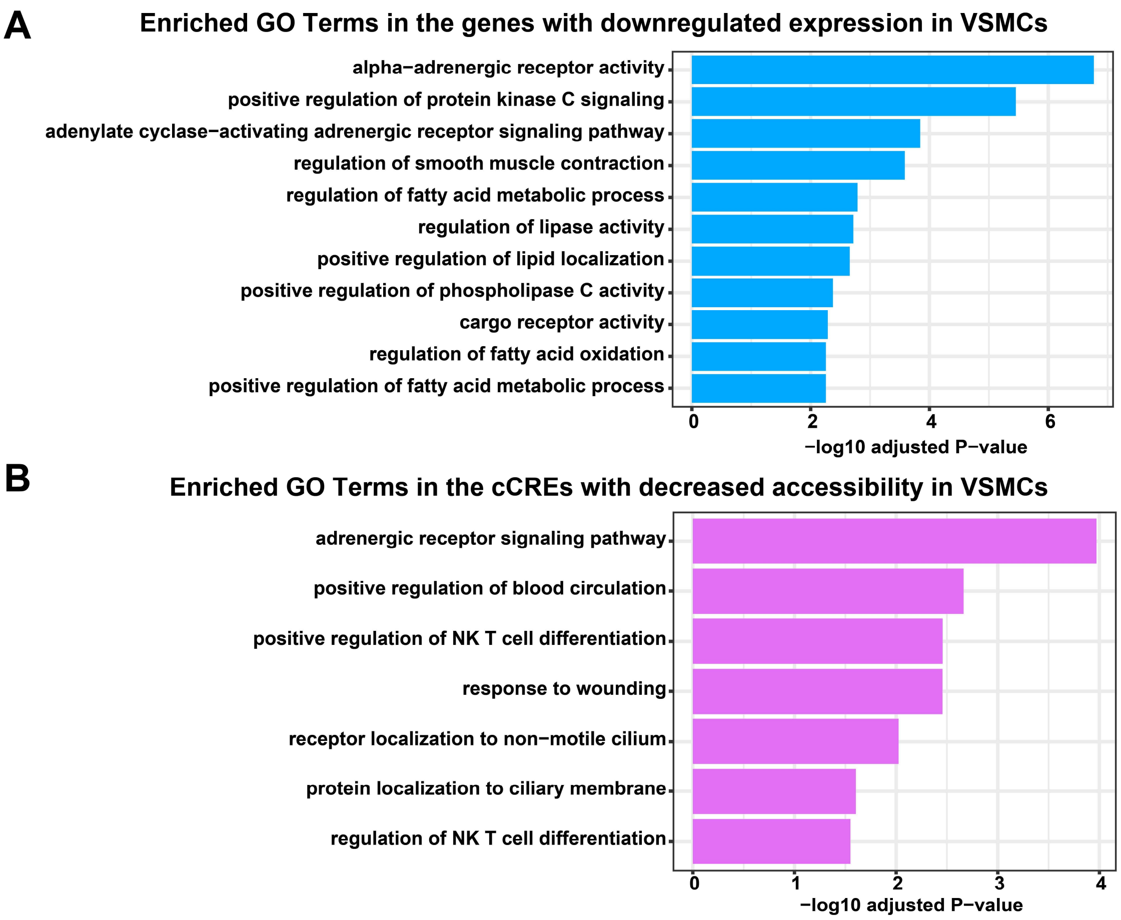


**Figure S9.** Enriched GO Terms in the downregulated genes and cCREs with decreased accessibility in VSMCS from MFS versus CTRL. A) Enriched GO terms enriched in the downregulated genes. The significance threshold was set to a Bonferroni-corrected $p$-value $< 0.05$. The hypergeometric test implemented in ClueGO was used. B) Enriched GO terms enriched in the cCREs with decreased accessibility. The significance threshold was set to a Bonferroni-corrected $p$-value $< 0.05$. The hypergeometric test implemented in GREAT was used.

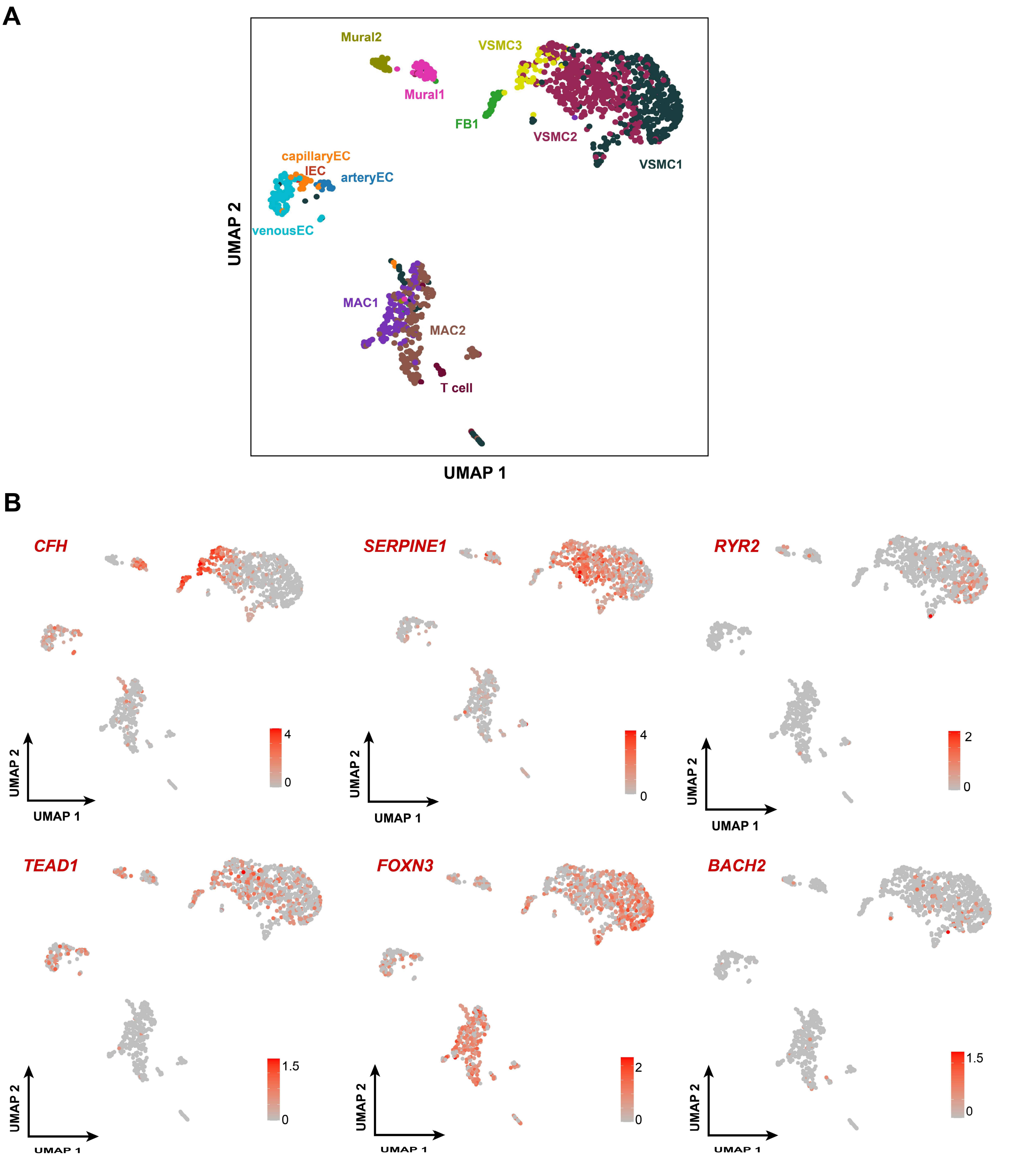


**Figure S10.** Reanalysis of a publicly available scRNA-seq dataset of aortic root tissue from one MFS patient (GEO sample accession ID: GSM4646673) and cellular annotation with our single-nucleus expression data as a reference. A) UMAP plot of the publicly available dataset showing that VSMC3 was located close to VSMC2, reflecting a special state of the phenotypic spectrum of VSMCs rather than fibroblasts. The cells are color-coded by subcluster. B) UMAP plots showing the expression of key markers and regulators. The annotation was performed using the label transfer workflow of Seurat.

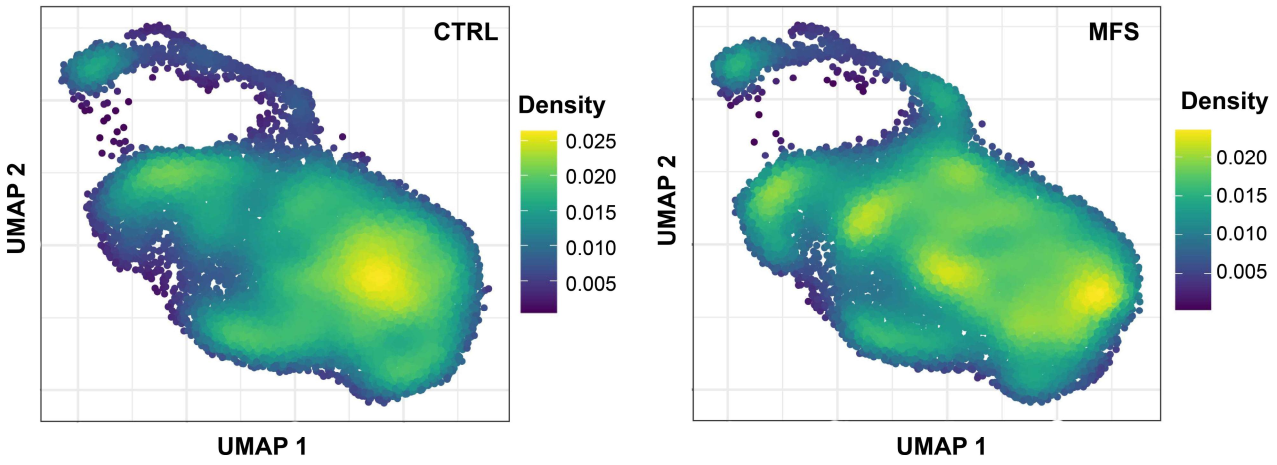


**Figure S11.** UMAP plots showing the nucleus density of VSMCs of the CTRL or MFS group.

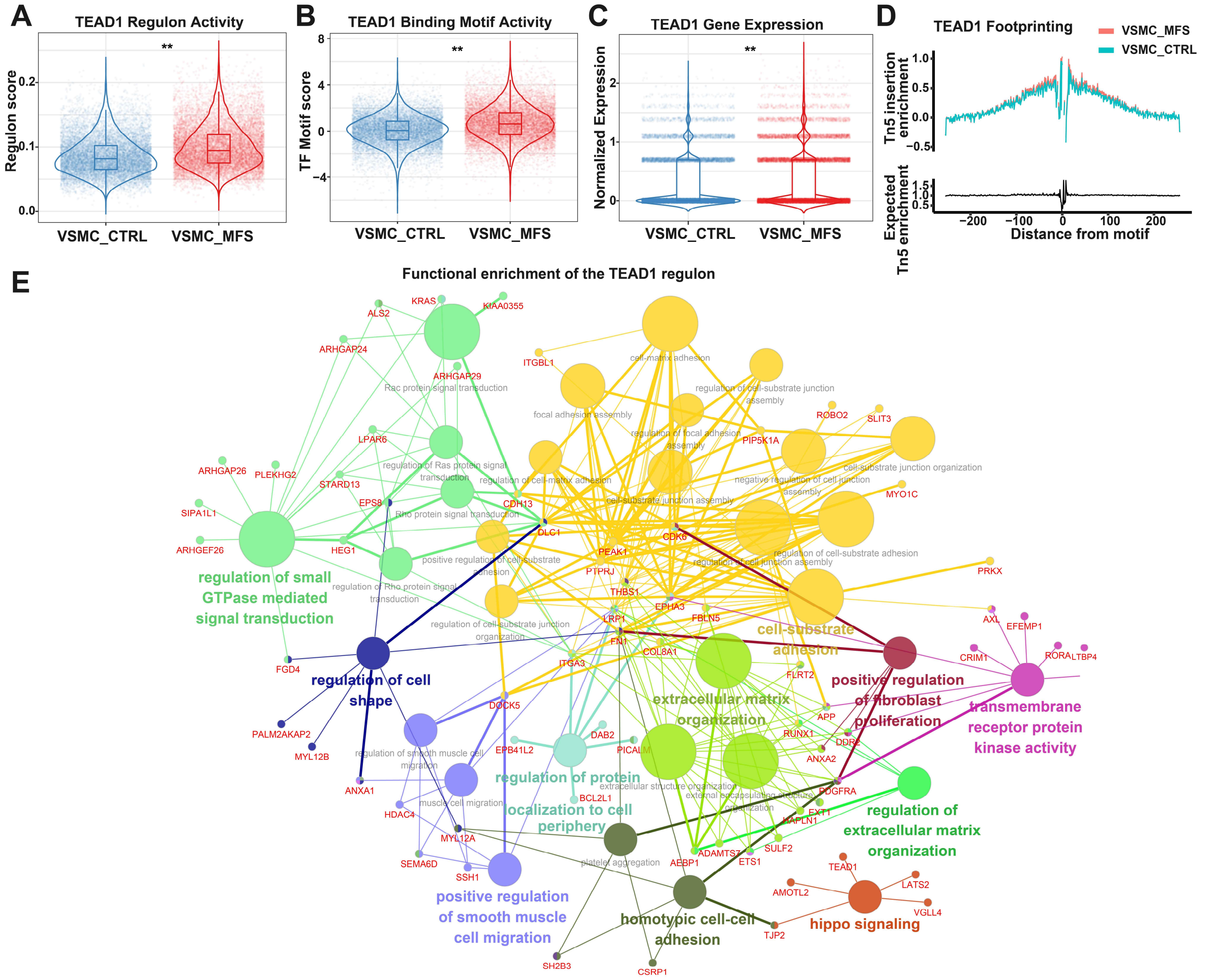


**Figure S12.** TEAD1 may function as a potential key regulator involved in the phenotypic modulation of aortic root VSMCs in MFS. A) Increased TEAD1 regulon activity in VSMCs of the MFS group versus the CTRL group. B) Increased TEAD1 binding motif activity in VSMCs of the MFS group versus the CTRL group. C) Increased gene expression of *TEAD1* in VSMCs of the MFS group versus the CTRL group. In A-C, **: *p*-

value < 0.05, Two-tailed Wilcoxon rank-sum test. D) Enhanced TEAD1 footprinting in VSMCs of the MFS group versus the CTRL group. E) Functional enrichment of the TEAD1 regulon (*TEAD1* and its predicted targets). The significance threshold was set to an adjusted *p*-value < 0.05. Each node denotes an over-represented GO term. The size reflects the statistical significance of the term.

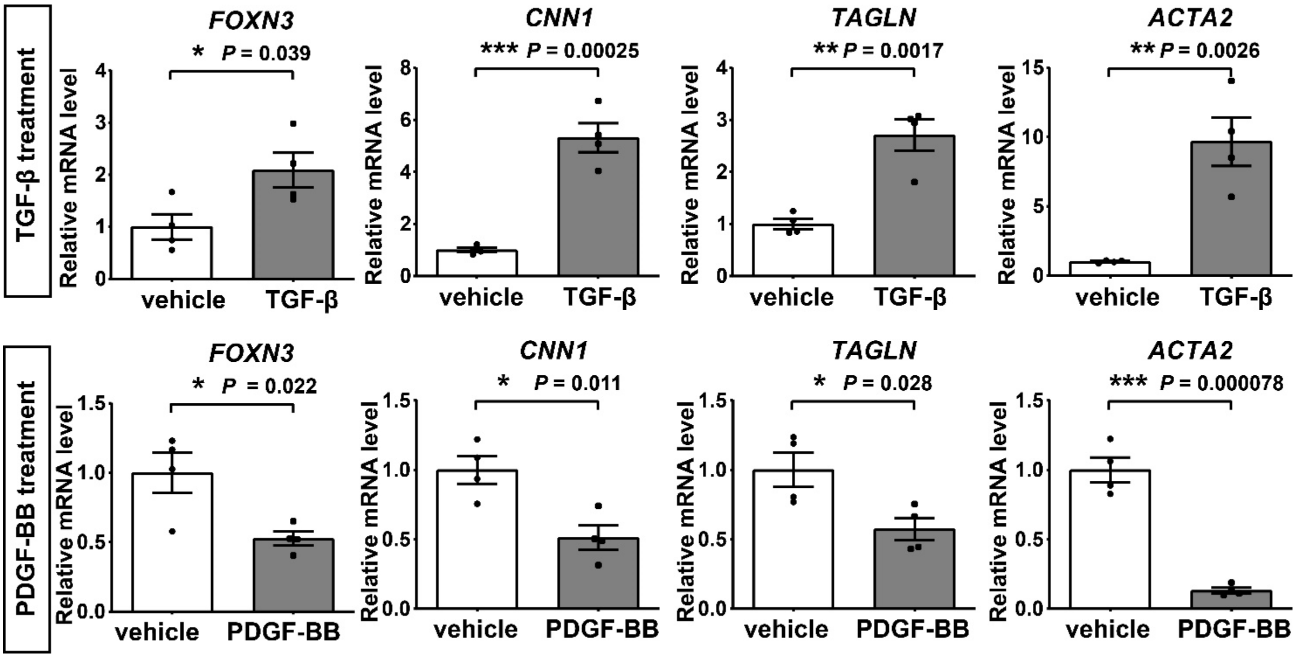


**Figure S13.** qRT-PCR results showing the mRNA expression changes of *FOXN3* and contractile marker genes in HASMCs following TGFβ or PDGF-BB stimulation. *: *p*-value < 0.05, **: *p*-value < 0.01, ***: *p*-value < 0.001, ns: not significant. Two-tailed Student's t-test. Data are presented as the mean ± SEM (n=4 independent experiments for each group).

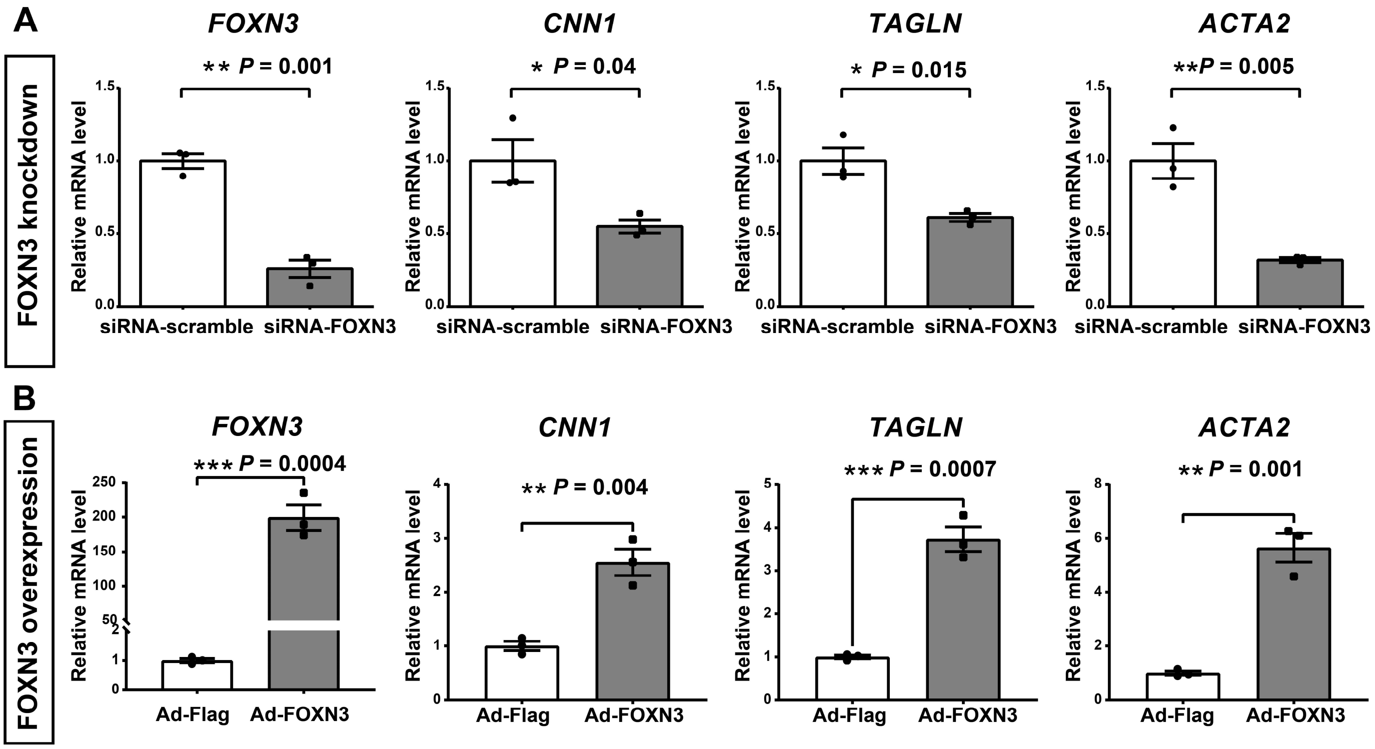


**Figure S14.** qRT-PCR results showing the mRNA expression changes of *FOXN3* and contractile marker genes in HASMCs following FOXN3 knockdown (A) or overexpression (B). *: $p$-value < 0.05, **: $p$-value < 0.01, ***: $p$-value < 0.001. Two-tailed Student's t-test. Data are presented as the mean ± SEM (n=3 independent experiments for each group).

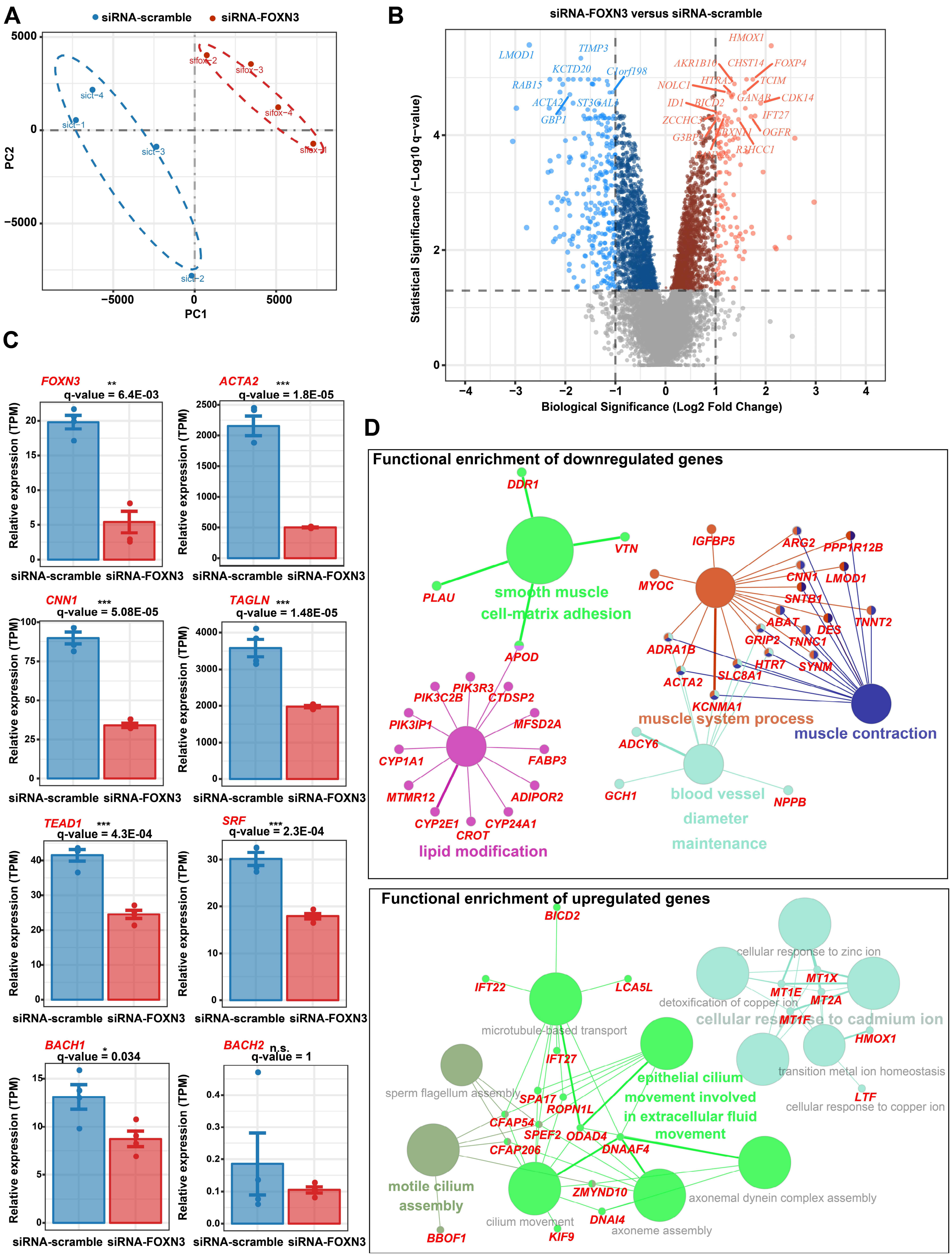


**Figure S15.** Bulk RNA-seq analysis revealed transcriptomic changes in HASMCs following siRNA-mediated knockdown of *FOXN3*. HASMCs were transfected with either scrambled siRNA (negative control) or

*FOXN3*-siRNA, and the bulk RNA-seq analysis was conducted 48 hours post-transfection. A) Principal component analysis revealed a clear discrepancy in the transcriptome between the knockdown group and the negative control group (n=4 independent experiments for each group). B) Volcano plot showing the differentially expressed genes detected in the knockdown group compared to the negative control group. The statistical significance threshold was set to a q-value < 0.05, and the biological significance threshold was set to an absolute log2 fold change > 1. The differential expression tests implemented in the R package sleuth was used. C) The expression of *FOXN3*, contractile marker genes *ACTA2* (α-SMA), *CNN1* (Calponin-1), and *TAGLN* (SM22α), and VSMC phenotypic modulation-associated regulators *TEAD1*, *BACH1*, *SRF*, *BACH2* following *FOXN3* knockdown. Data are presented as the mean ± SEM (n=4). D) Network plot showing functional enrichment of the downregulated genes (upper panel) and the upregulated genes (lower panel) in the knockdown group compared to the negative control group. The significance threshold for hypergeometric tests was set at a Bonferroni-corrected *p*-value of less than 0.05. The smaller the *p*-value, the larger the circle size.

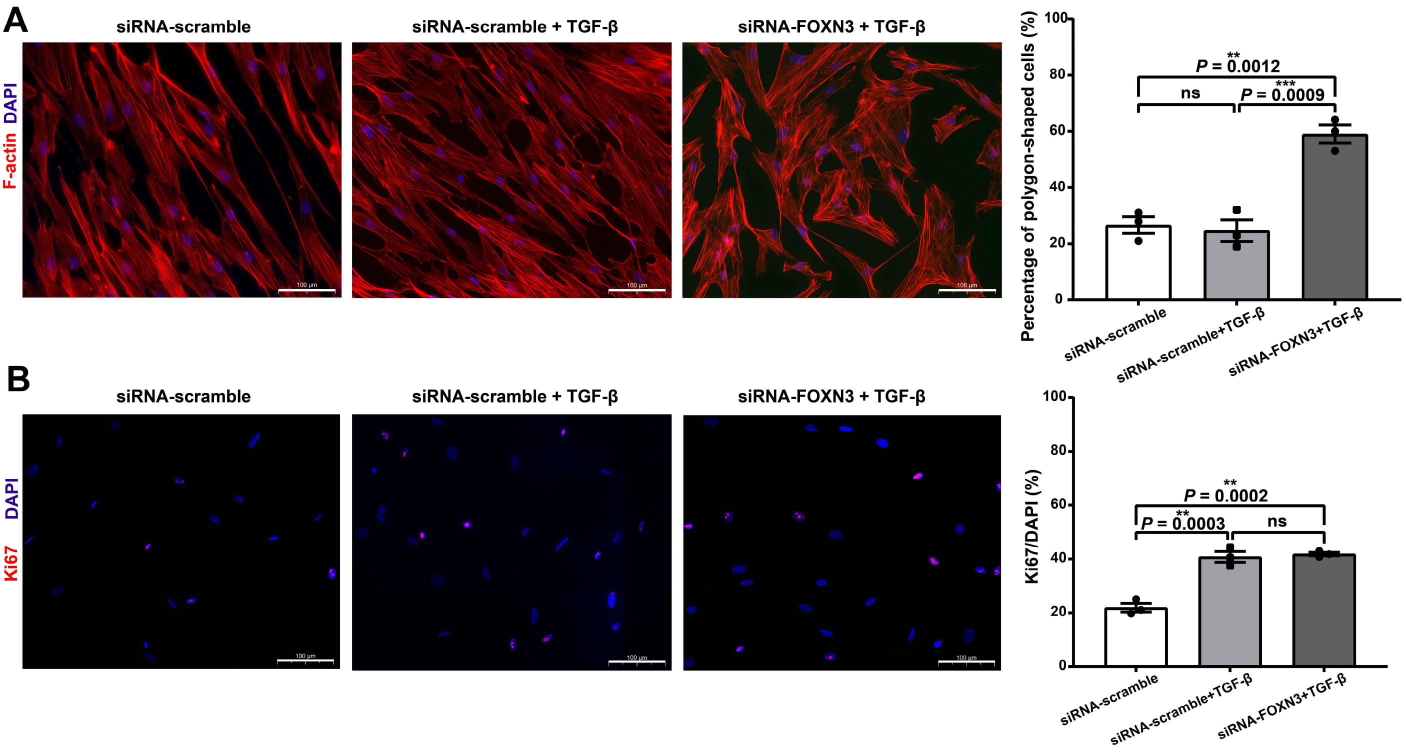


**Figure S16.** Effects of *FOXN3* knockdown on the morphology and proliferation of HASMCs in response to TGF-β treatment. A) Representative immunofluorescence staining images of F-actin (red) in HASMCs infected with scrambled siRNA (10 nmol/L) or FOXN3-siRNA (10 nmol/L) for 48 h and then subjected to TGF-β (10 ng/mL) treatment for 24 h. B) Representative immunofluorescence staining images of Ki-67 (red) in HASMCs infected with scrambled siRNA or FOXN3-siRNA and then subjected to TGF-β treatment. For all experiments, the data are presented as the mean ± SEM (n=3). One-way ANOVA followed by multiple comparisons using Tukey's method. *: *p*-value < 0.05, **: *p*-value < 0.01, ***: *p*-value < 0.001, ns: not significant. The percentage of polygonal-shaped cells or Ki-67-positive cells in each image was calculated as the mean of the measurements in at least five representative views. Nuclei stained by DAPI are indicated in blue. Scale bar: 100 μm.

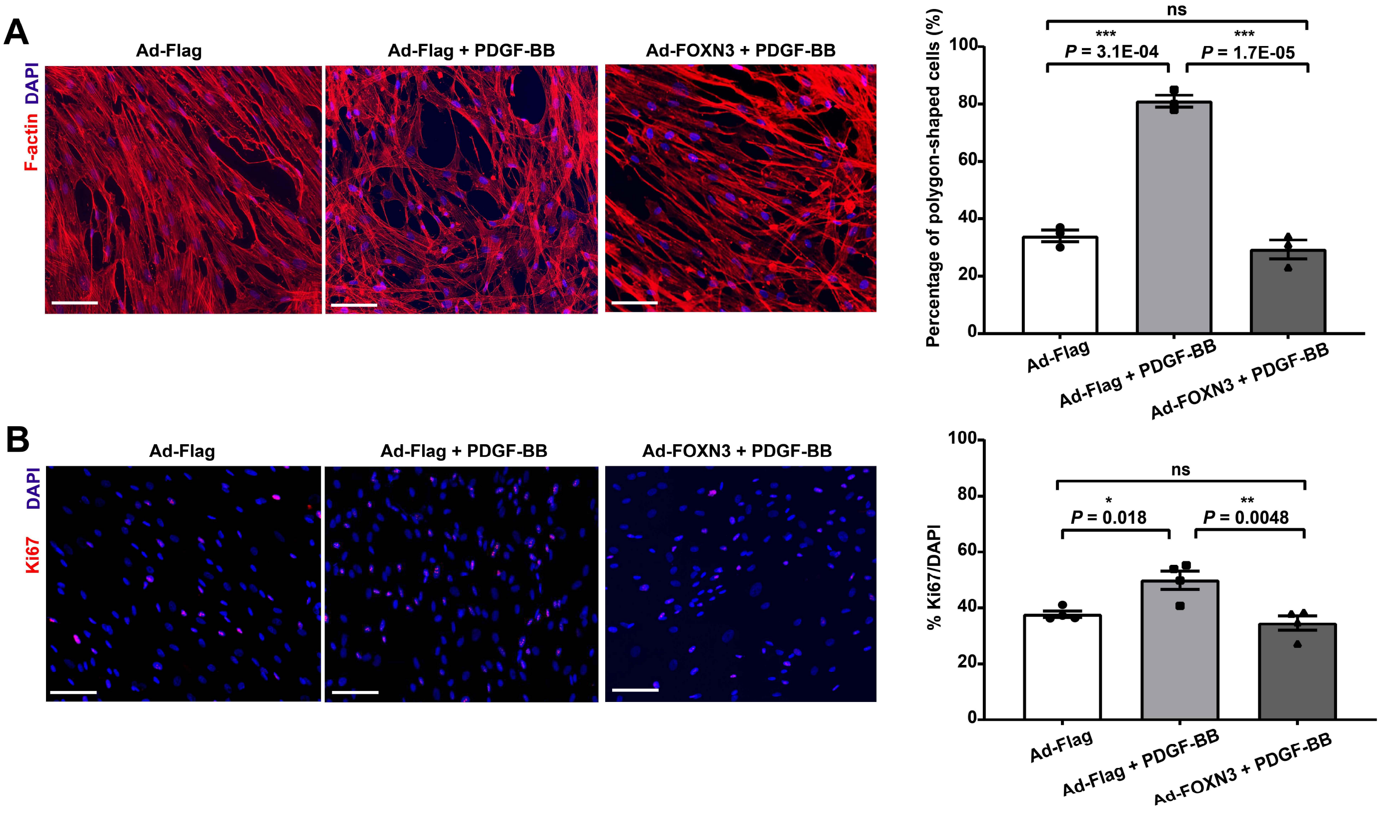


**Figure S17.** Overexpression of *FOXN3* attenuated the phenotypic modulation and proliferation of HASMCs induced by PDGF-BB stimulation. A) Representative immunofluorescence staining images of F-actin (red) in HASMCs infected with Ad-Flag or Ad-FOXN3 for 72 h and then subjected to PDGF-BB (20 ng/mL) treatment for 24 h. B) Representative immunofluorescence staining images of Ki-67 (red) in HASMCs infected with Ad-Flag or Ad-FOXN3 and then subjected to PDGF-BB treatment. For all experiments, the data are presented as the mean ± SEM (n=3). One-way ANOVA followed by multiple comparisons using Tukey's method. *: $p$-value < 0.05, **: $p$-value < 0.01, ***: $p$-value < 0.001, ns: not significant. The percentage of polygonal-shaped cells or Ki-67-positive cells in each image was calculated as the mean of the measurements in at least five representative views. Nuclei stained by DAPI are indicated in blue. Scale bar: 100 μm.
